# Extending the application of the SCA/Sectors method for the identification of domain boundaries and subtype specific residues in multi-domain biosynthetic proteins: Application to Polyketide Synthases

**DOI:** 10.1101/2021.06.30.450545

**Authors:** Tuğçe Oruç, Christopher M. Thomas, Peter J. Winn

**Affiliations:** School of Biosciences, University of Birmingham, Edgbaston, Birmingham, B15 2TT, United Kingdom; The Institute of Microbiology and Infection, University of Birmingham, Edgbaston, Birmingham, B15 2TT, United Kingdom; Centre for Computational Biology, Haworth Building (Y2), University of Birmingham, Edgbaston, Birmingham, B15 2TT, United Kingdom

## Abstract

Polypeptides with multiple enzyme domains, such as type I polyketide synthases, produce chemically complex compounds that are difficult to produce via conventional chemical synthesis and are often pharmaceutically or otherwise commercially valuable. Engineering polyketide synthases, via domain swapping and/or site directed mutagenesis, in order to generate novel polyketides, has tended to produce either low yields of product or no product at all. The success of such experiments may be limited by our inability to predict the key functional residues and boundaries of protein domains. Computational tools to identify the boundaries and the residues determining the substrate specificity of domains could reduce the trial and error involved in engineering multi-domain proteins. In this study we use statistical coupling analysis to identify networks of co-evolving residues in type I polyketide synthases, thereby predicting domain boundaries. We extend the method to predicting key residues for enzyme substrate specificity. We introduce bootstrapping calculations to test the relationship between sequence length and the number of sequences needed for a robust analysis. Our results show no simple predictor of the number of sequences needed for an analysis, which can be as few as a hundred and as many as a few thousand. We find that polyketide synthases contain multiple networks of co-substituting residues: some are intradomain but most multiple domains. Some networks of coupled residues correlate with specific functions such as the substrate specificity of the acyl transferase domain, the stereo chemistry of the ketoreductase domain, or domain boundaries that are consistent with experimental data. Our extension of the method provides a ranking of the likely importance of these residues to enzyme substrate specificity, allowing us to propose residues for further mutagenesis work. We conclude that analysis of co-evolving networks of residues is likely to be an important tool for re-engineering multi-domain proteins.

**Author summary:** Many important compounds such as antibiotics or food flavourings are produced naturally by molecular factories within plant, fungal and bacterial cells. These molecular factories typically comprise a complex of multiple interacting enzymes, each enzyme being a stage in a molecular production line. Often the enzymes are connected together as subsections of the same amino acid chain, i.e. protein, with the amino acid chain folding into the separate functional enzymatic domains that comprise the production line. Polyketide synthases are such multi-domain proteins, and their products often have antibacterial, antifungal and antitumoric effects. Engineering polyketide synthases thus has the potential to produce novel drug candidates. We applied and developed statistical approaches to detect where in an amino acid sequence the boundaries are between different domains, potentially allowing these regions to be swapped around for the synthesis of novel compounds. We used the same approaches to identify parts of the amino acid chain important for the function of different types of domain, pointing to how they might be modified to make novel compounds. These analyses agree with published experimental data and allow us to make novel predictions, which we expect to help experimentalists produce novel compounds of commercial and pharmaceutical interest.

## 1 Introduction

Polyketide synthases (PKSs) are protein complexes that produce polyketides which constitute a large class of secondary metabolites. Polyketides have a wide range of pharmacological properties, including antibiotic, antitumor, and antifungal activity. PKSs come in three types but our focus here is on Type I modular PKSs, which are characteristically megapolypeptides consisting of domains with specific enzymic functions covalently joined by flexible linker regions, and which are responsible for almost one-third of commercial pharmaceuticals [1].

Type I modular PKSs consist of consecutive modules, each module typically elongating the polyketide chain and passing the resulting intermediate chain to the following module. A minimal elongating module consists of ketosynthase (KS), acyltransferase (AT) and acyl carrier protein (ACP) domains, which collaborate to elongate the polyketide chain. A minimal loading module, the first module of the biosynthetic pathway, consists of only the AT and ACP domains, the initial substrate being captured by the AT domain and transferred to the ACP domain for transfer to, and elongation by, downstream modules (Fig 1A, B). In the elongation process, the KS domain performs a Claisen condensation reaction, which ligates an extender unit, initially bound to the ACP of the elongating module, onto the elongating chain bound to the KS, forming a carbon-carbon bond and leaving a keto group at the beta carbon (Fig 1A, B). The resulting elongated product is bound to the ACP and may then be passed to the next downstream KS for further elongation or, at the end of the biosynthetic pathway, released by a thioesterase (TE) domain.

**Fig 1.**
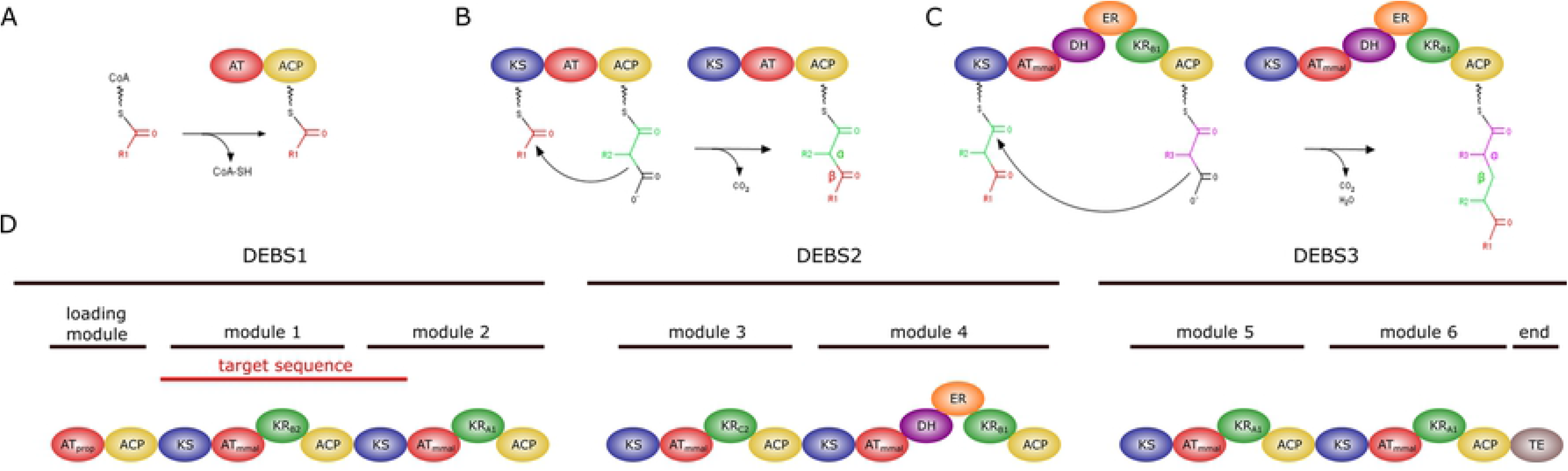
Schematic illustration of polyketide synthesis showing how different domain compositions and domain subtypes lead to diversity in polyketide structure. (A) The initial substrate (starter unit) is loaded onto a PKS via a loading module. (B) A minimal PKS includes the KS, AT and ACP domains, the AT loading an extender unit onto the ACP, which then presents it to the substrate bound to the KS, elongating that substrate and resulting in a keto group at the *β*-carbon. (C) One or more further rounds of elongation can occur, and the presence of modification domains (KR, DH, ER) reduces the keto group at the *β*-carbon, shown by a *β* on the figure, which results in a fully reduced carbon chain when KR, DH and ER domains are all present in the module. (D) The modular organization of the DEBS system, which consists of three polypeptides (DEBS1, DEBS2, DEBS3) and six elongation modules. The fragment of DEBS1 used as the basis of our analysis is marked in red and has a KS at the N- and C-termini of the fragment (KS1 and KS2, respectively). This sequence was chosen since we would expect it to form a structurally closed unit, since the PKSs are actually homodimers with KS1 of one PKS polypeptide binding to KS1 of another polypeptide chain and KS2 similarly binding another KS2, effectively bounding the structure of the module [3, 4]. Choosing this sequence for study also fits the most prevalent definition of a PKS module, with the KS at the N-terminus as show in the figure, but also for the possibility that the more natural definition of a module might be with the KS at the C-terminus rather than the N-terminus [5, 6]. Although every module of DEBS includes a KR domain, they are of different subtypes, A1, B2, C2, which are described later in the text.

The AT domain of the elongating module captures the extender unit and loads it onto the ACP domain of that module. Although there are several types of extender unit, AT domains specific for methylmalonyl-CoA or malonyl-CoA are the most abundant ones [2]. Some AT domains select only one type of extender unit, but others can accept multiple types. In some PKS systems, the AT domain is present within the module and these systems are called *cis*-AT systems. On the other hand, an AT domain is not found within the modules of all PKSs. In some systems the AT is a discrete protein that acts in *trans* so these PKSs are called *trans*-AT systems.

In some modules, in addition to the KS, AT and ACP domains, ketoreductase (KR), dehydratase (DH) and enoylreductase (ER) domains exist, which provide modifications to the elongating chain (Fig 1C). The KR domain reduces the keto group at the *β* carbon, which was left by the Claisen condensation, to a hydroxyl group. The DH domain removes the hydroxyl group leaving a double bond and the ER domain reduces the double bond into a single bond. Depending on the exact composition of a PKS module the beta carbon may thus be left as a keto, a hydroxyl, an enoyl or a methylene group. Further diversity in the structure of the elongating chain also arises due to the subtypes of the modification domains. For example KR domains have six subtypes (A1, A2, B1, B2, C1 and C2) which result in different stereochemistry in the product [7] (Fig 1D). Type A KR domains produce a hydroxyl group with L-configuration at the beta-carbon, whereas type B KR domains produce a D-configured hyrdoxyl group. Types A and B can be further classified by their ability to epimerise the alpha carbon. A1/B1 leave a D-configured alpha carbon, thought to be a result of the stereochemistry of the extender unit [8], whereas A2/B2 leave an L-configured alpha carbon, a result of enzyme catalysed epimerisation [9]. Type C1 have no catalytic ability, whereas type C2 only epimerise the alpha carbon to be L-configured [9].

New polyketide variants can be produced by modifying the extender unit specificity of the AT domain, which will change the moiety added to the growing polyketide chain, or by changing the KR domain subtype [7, 10–13], which can modify the extended polyketide as described above. One common approach to changing AT specificity is to swap the native domain with the corresponding domain from another PKS system or module. The challenging points of domain swapping experiments are understanding the compatibility for interaction between the inserted domain and the domains on the host system, and correctly identifying the functional boundaries of the domain, both of which are thought to be needed to maintain structural and functional integrity. Although studies of the AT domain swapping have been going on for more than two decades, successful swapping was achieved only recently [14–19]. Experimenting with different domain boundaries, swapping the AT domain from EPOS module 4 into the position of the AT in DEBS module 6 identified two functional variants from four constructs [19]. The same work demonstrated the functionality of these domain boundaries in other systems. On the other hand, no similar study has been published regarding KR domain boundaries, to the best of our knowledge. KR swap experiments have been published but their success varies with KR domain subtype [20–23].

Another approach to engineering polyketide variants has been point mutation of residues that are thought to be critical for the functional specificity of the domain. For example, sequence analysis revealed that methylmalonyl specific AT domains bear a YASH motif whereas the corresponding position in the sequence of malonyl specific AT domains is a HAFH motif. However, switching extender unit specificity by mutating only the YASH/HAFH motifs led to incorporation of both types of extender unit, indicating that specificity is not solely defined by these motifs [24–26]. This motivated the search for additional residues that are functional in extender unit binding. A particularly successful attempt studied malonly/methylmalonyl/ethylmalonyl specificity of three ATs in the salinomycin producing pathway [27]. Molecular docking and simulation identified substrate binding residues in addition to the YASH/HAFH motifs and mutating these produced AT domains with altered extender unit preferences. Although the modified domains loaded substrate with similar efficiency to the wild type ATs they had *K_m_* for the cognate holo-ACP that was up to an order of magnitude less favourable than the wild type interactions. Thus, although the challenge of loading an AT with an alternative substrate seems to have been addressed, the challenge of mutating AT specificity in the context of the PKS still needs more work. These experiments showed that switching the extender unit specificity from one type to another is possible only if all the key substrate binding residues are identified with further work presumably needed to characterise the residues that couple an AT’s function to its cognate ACP.

Similarly, site-directed mutagenesis has been applied to KR domains to switch the KR subtypes. In type-B KR domains, approximately 57 residues before the catalytic tyrosine there is a highly conserved LDD motif (the second aspartic acid residue is strictly conserved). This motif is absent in type-A KR domains, but these conserve a tryptophan eight residues before the catalytic tyrosine. Although mutating these critical residues has given some promising results, with type-A1 and type-B2 being mutated to type-A2, switches to other types have not yet given high product yields [28, 29].

Overall, although these studies provided better understanding of the domains and their subtypes, a complete conversion from one subtype to another, whilst maintaining high product yields, has yet to be achieved *in vivo*. However, the current results make it clear that identifying what constitutes a functional domain is critical for the success of domain swapping experiments. Furthermore, although sequence analysis has provided fingerprint motifs of the AT and KR subtypes, the low levels of success in experiments that switch such motifs suggests a need to identify additional residues necessary for the specific functions of these domains. Thus if residues that are closely functionally coupled could be easily identified this would expedite the problem of engineering novel polyketide synthase pathways. By closely functionally coupled we mean either residues that work together structurally as an enzymic domain or as the group of key residues responsible for subtype specificity.

Many computational approaches have been developed to detect co-substituting residue pairs or groups [30–33] and thence to identify networks of residues that appear to work together [33]. These methods are predicated on the process of evolution leading to coupled amino acid changes. To understand how this might happen consider the following: if a mutation occurs at a position in the three dimensional structure of a protein it may lead to suboptimal protein function, possibly leading to the death of the host organism and thus to the loss of the mutation from the gene pool, unless one or more further mutations compensate for that first change. Amino acid substitution patterns within multiple sequence alignments evidence covariance between positions in the alignment, supporting this idea of coupled amino acid substitutions, and analysis of this covariance can identify functionally and structurally coupled amino acids.

One approach to identifying networks of coevolving residues is sectors analysis [33, 34]. Sectors analysis is built on the statistical coupling analysis (SCA) of Ranganathan and co-workers and depends on having the sequences of a family of related proteins that are sufficiently divergent that a significant proportion of the possible substitutions that can occur have occurred in at least one member of the sequence family. SCA creates a matrix that represents the covariance of amino acids between the different columns of a multiple sequence alignment. Each entry in the covariance matrix is weighted by a measure of the amino acid distribution (the derivative of the Kulback-Leibler divergence with respect to amino acid frequency) in the pair of sequence positions under consideration [34]. This upweights the contribution of positions with residue types that deviate from a precalculated average background frequency. Conservation of rare residue types such as cysteine and tryptophan thus have a strong upweighting effect, whereas the absence of a common residue type such as leucine will also have an upweighting effect, as will their conservation, although this will not be as great as the upweighting that a highly conserved tryptophan receives. The covariance matrix is then decomposed via eigenvalue spectral decomposition and independent component (IC) analysis, to identify networks of residues that are particularly strongly coupled with regard to their tendency to co-vary. Since they coevolved, residues in each group (sector or IC) are expected to be functional for a specific purpose.

The idea of functionally independent sectors was first validated in the serine protease enzyme family. Computational analysis identified three almost independently co-varying sectors, two of which were confirmed with experiments in the laboratory. Mutational analysis showed that one sector seemed to control the stability of the serine protease fold but had little effect on catalytic efficacy, and another sector was important for catalysis but had little effect on protein stability [33]. This approach has since been applied to several protein families including G-protein coupled receptors, DHFR, *β*-lactamase and the pancreatic-type ribonuclease superfamily [34, 35]. Although sectors analysis has been successfully applied on single-domain systems, there is no study to date that has analysed a multidomain protein.

The polyketide synthases provide a good system for testing how sectors analysis performs on a multidomain protein and sectors analysis has the potential to identify networks of residues defining domain boundaries or critical residues for enzyme specificity, and is likely to be complementary to existing methods (e.g. [36, 37]). The DEBS polyketide synthases are responsible for the production of the antibiotic erythromycin and have been investigated extensively and thus have a rich literature from which to draw experimental data to validate a sectors analysis. We analysed proteins related to the first elongating module of the DEBS pathway, including the KSs at the N- and C- termini of the module, which has domain composition KS1_AT_KR_ACP_KS2 (Fig 1D). We detect approximately 22 ICs representing coevolved residue groups, some of which correlate with known domain boundaries, others with subtypes of the PKS domains. Some ICs span multiple domains suggesting that ICs can give further insight into the interaction pattern of the residues within and between domains and inter-domain linkers. We perform bootstrapping analysis to test the robustness of our IC predictions and to provide recommendations for the number of sequences required for sectors analysis. To the best of our knowledge, this is the first study that applies sectors analysis to multidomain proteins like the PKSs.

## 2 Results and Discussion

### 2.1 Determining the number of sequences needed for a sectors analysis

The literature suggests that 100 sequences are sufficient for a sectors analysis [34], based on calculations from one example system. In contrast, other covariance methods such as direct coupling analysis require thousands of sequences to give reliable results. Moreover, the DEBS module that we wish to analyse has 1901 amino acids, which is considerably larger than any sequence previously analysed by sectors analysis [34, 35] and which we thus anticipated might need a larger number of sequences to obtain robust results. We, therefore, analysed several uni-domain proteins to see whether we could detect a trend between the length of a protein and the number of sequences adequate for a sectors analysis.

To test how the number of sequences in an MSA affected the ICs, varying numbers of sequences were randomly selected from a source MSA, without replacement, and ICs were determined. This was repeated three times for each different number of sequences (n=3). To see the similarity between the ICs from subsampled MSAs and the ICs from the source MSA a similarity score (ss) was calculated between the sets of ICs, ranging from 0 (no similarity) to 1 (every IC calculated for the first MSA can be paired with an IC calculated for the second MSA with at least 70% identity).

For most of the proteins analysed, 100 sequences in the MSA is not sufficient to reproduce ICs similar to the source MSA ICs, irrespective of the length of the protein (S2 Fig). On the other hand, the similarity score converged to one once the MSA had 90% (but often fewer) of the effective sequences found in the original MSA, where an effective sequence is the number of sequences in the alignment whose pairwise sequence identity compared to every other sequence is 80% or less.

However, no trend was detected between the length of the protein and the effective number of proteins in the MSA at which the ICs of the full MSA were recovered (the saturation point, 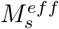, Fig 2 and S2 Fig). Nor was there a trend with respect to the saturation ratio, the ratio of 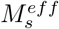 to *M^eff^*, the effective number of sequences in the full alignment (Fig 2).

**Fig 2.**
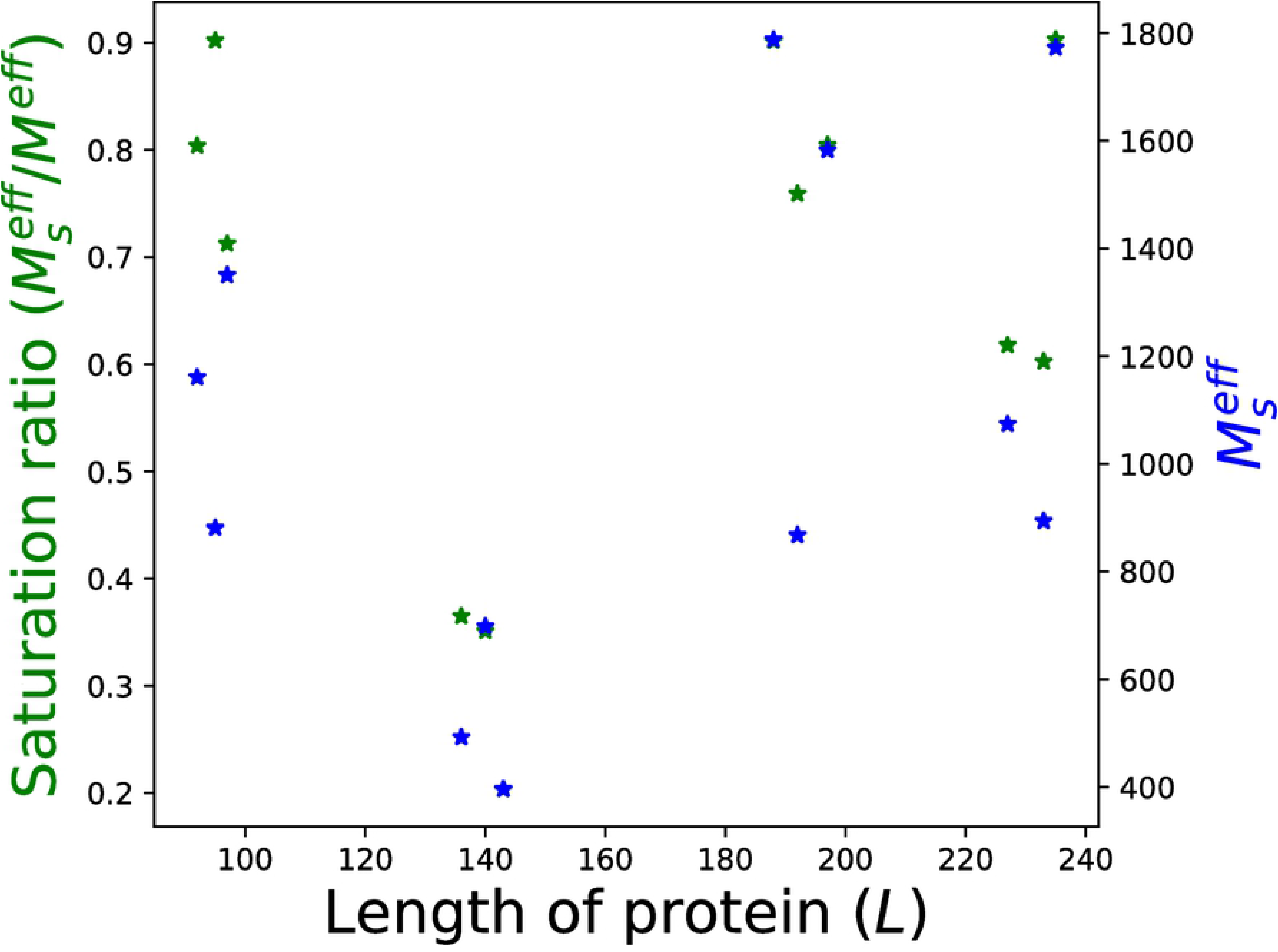
A trend could not be detected between the length of the proteins (*L*) and the saturation ratios 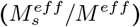 or 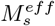 values. 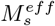 values are defined as the minimum number of sequences adequate to achieve the same set of ICs as the source MSA (with *M^eff^* number of sequences). Selected 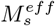 values for each protein are shown in S2 Fig.

For most proteins, the subsampled MSAs with the fewest sequences did not reproduce the ICs identified for the full MSA, but some ICs were found in all subsamplings for each protein (S3 Fig). Consistently detecting an IC does not seem to depend on the number of residues in the IC (S4 Fig) and although the number of sequences in the MSA is critical for the sector analysis, some ICs can be successfully detected even when the MSA has few sequences. This suggests that, although the number of sequences in the MSA is criticial for the sector analysis, some ICs can be successfully detected even when the MSA is not comprehensive enough. On this basis the subsequent analysis has focused only on ICs that appear to be robust in the sense that they can be detected consistently in subsets of the sampled proteins as well as the whole set.

### 2.2 Twenty Two Independent Components were Robustly Identified in a sequence alignment of PKS modules with sequence similarity to DEBS Elongation Module One

To identify networks of co-evolving residue positions within a module of a Type I PKS, an initial multiple sequence alignment (MSA)was generated from protein sequences identified by HHblits, using as the query term the sequence of the first elongating module of the DEBS system plus the KS domain of the second module, KS1_AT_KR_ACP_KS2 (S1 Fig). After processing the MSA to remove highly gapped positions and sequences, and highly identical ones, as described in Methods, 2245 sequences remained in the alignment. Only 682 of the sequences have the exact domain composition of the input sequence. The remaining sequences have either a missing domain (like AT or KR) or an additional domain (like DH or ER). Having 600 or fewer sequences in the alignment was sufficient for convergence of the sector analysis for some of the proteins we analysed in the previous section, but here the input sequence is longer than these proteins. Thus we investigated the effect of the number of sequences in the alignment on the ICs.

The similarity between the ICs derived from a subsampled alignment and that of the full alignment of 682 sequences increases as the number of sequences in the subsample increases. The similarity does not converge to a point where the addition of new sequences has no further effect (S5 Fig). Indeed it is never larger than 0.5 similarity, averaged across all ICs. This indicates that the alignment may be too small to robustly identify the ICs. Therefore, we included all the 2245 sequences we had obtained from the preliminary sequence search. Although all sequences had at least two KS domains and an ACP, sequences were also included without AT and/or KR domains or with DH and/or ER domains.

Although the inclusion of additional sequences was not adequate to obtain the convergence of the results, the maximum similarity score is higher than the one where only sequences with KS1_AT_KR_ACP_KS2 domain composition were included in the MSA (S6 Fig). As noted in detail later, the conclusions arising from our analysis were tested to ensure that the additional sequences did not lead to misleading artefacts.

Analysis of the full sequence alignment gave 34 ICs with two or more residues. We performed resampling by randomly drawing from the full MSA three lots each of 100, 500, 900, 1300, 1700 and 2100 sequences. Some ICs were detected even when there were only a few sequences in the alignment, but some occurred three or fewer times, over all samplings, and these were discarded as not being robustly defined. IC 25 was also discarded as it consisted of only two residues. After discarding the aforementioned ICs this left 22 ICs(Figs S7 Fig, S8 Fig and Fig 3).

**Fig 3.**
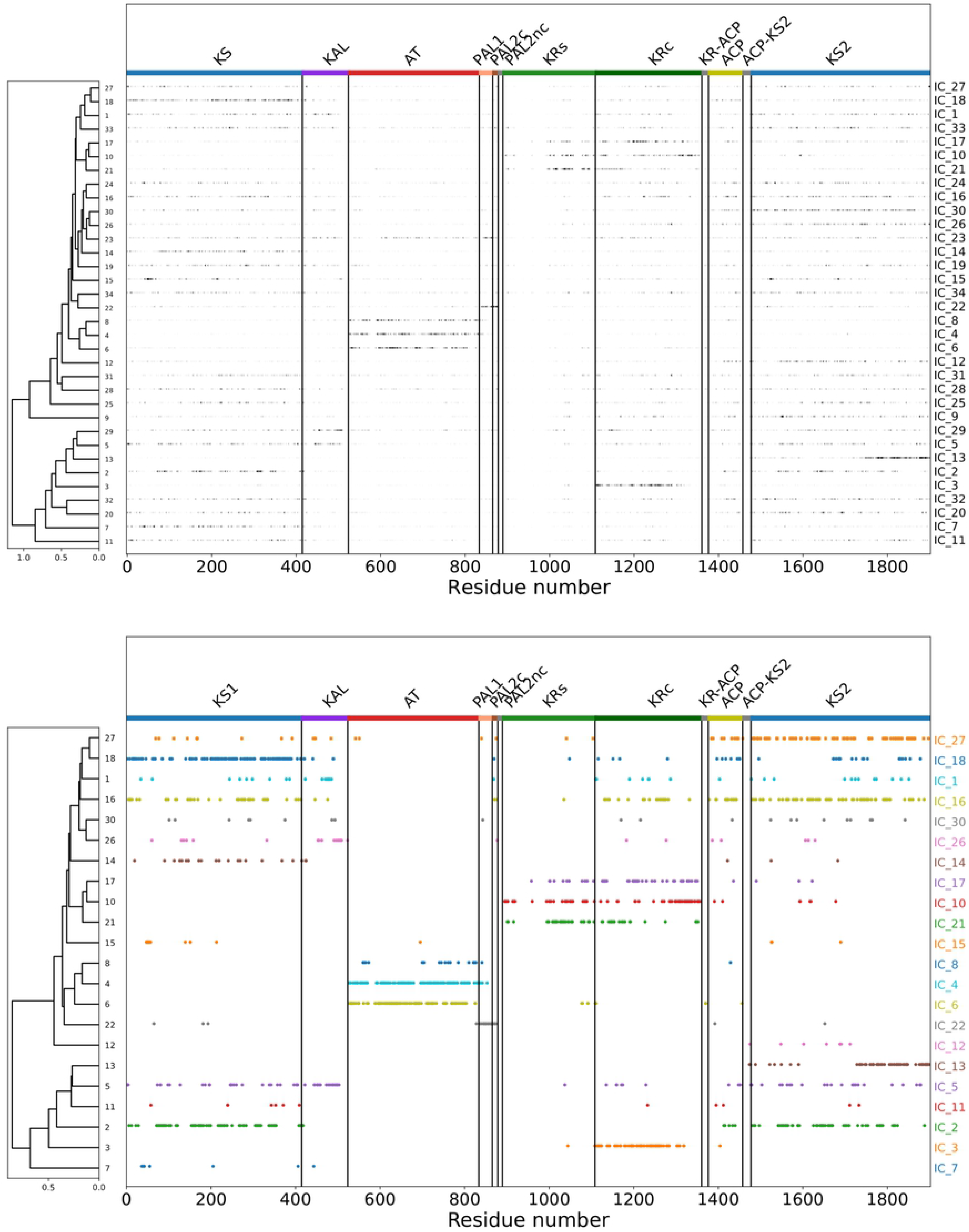
SCA analysis revealed 34 coevolved residue groups of which 22 were consistent in subsamples of sequences. The MSA of sequences found by an hhblits search with the DEBS module 1, with domain architecture KS1_AT_KR_ACP_KS2, was analysed by SCA and independent component analysis. The left side of each panel shows the hierachical clustering of ICs, as described in Methods, with the right side showing how individual residues contribute to each IC. The upper panel shows all ICs from the full sequence analysis and has residues shaded darker the greater their contribution to the independent component, which can be interpretted as the strength of coupling a residue has to the co-evolving residue network of the IC. The lower panel shows the ICs found in multiple subsamples of the full sequence alignment, with residues assigned to an IC according to the distribution of residue scores along that IC (based on p=0.95 threshold). KS1: Ketosynthase of module 1 (residue 1 to 414), KAL: KS-AT linker (residue 415 to 523), AT: Acyltransferase residue (524 to 833), PAL1: post-AT linker 1 (residue 834 to 865), PAL2c: post-AT linker 2 conserved region (residue 866 to 878), PAL2nc: post-AT linker 2 non-conserved region (residue 879 to 889), KRs: Ketoreductase structural integrity region (residue 890 to 1108), KRc: Ketoreductase catalytic region (residue 1109 to 1360), ACP: acyl carrier protein (residue 1377 to 1457), KS2: Ketosynthase of module 2 (residue 1478 to 1901). Residue 1 here corresponds to residue 505 of the polypeptide chain of DEBS1 since we have ignored the loading domains of DEBS1, which acquire and activate the starter unit for biosynthesis

### 2.3 Functional domain boundaries can be detected

Some independent components consist of residues that are predominantly from only one domain but most ICs have signal from multiple domains and linkers (Fig 3). However, even where an IC spans multiple domains, the strongest couplings are sometimes still confined to one domain. Further insight into the coevolutionary relationships within a protein can also be gained by looking at residues coupled across two ICs. Although an IC is a grouping of residues that coevolve together, predominantly independently of other ICs, there can nonetheless be couplings between residues in different ICs (S9 Fig). For this reason, previous work has grouped together ICs with high inter-IC coupling, each grouping termed a sector [34]. However, there is no standard way to group ICs into sectors. Here, we therefore do not definitively define sectors but instead make hierarchical clusterings of ICs based on the average interaction score between residues in each pair of ICs as explained further in Methods section 4.6 (Fig 4). Such hierarchical clustering shows that AT and KR domains are characterised by several ICs that cross-couple (Fig 3) and which have residues predominantly from the same domain, thus allowing us to define domain boundaries.

**Fig 4.**
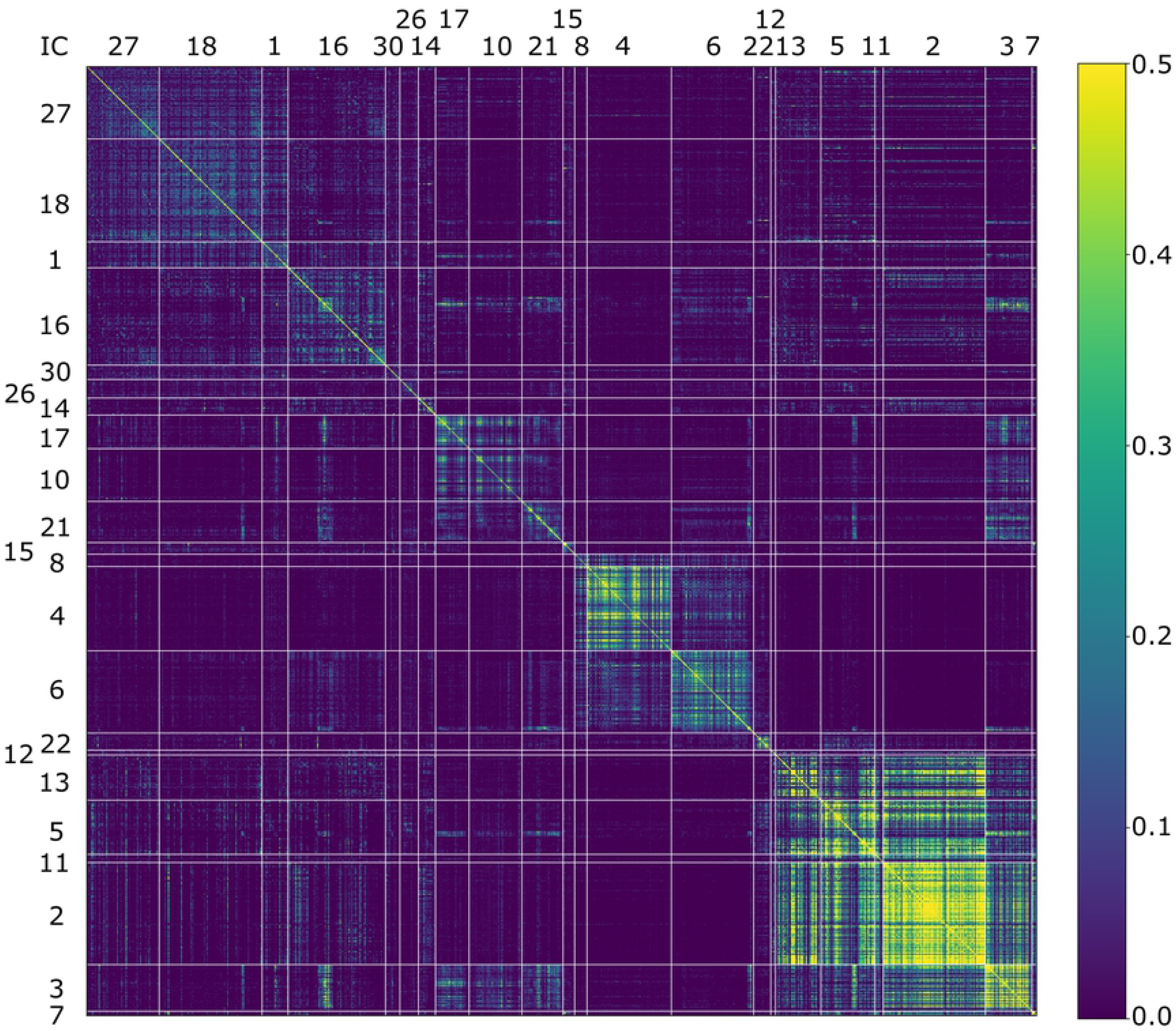
Cross-coupling between residues in the robust ICs from the MSA of sequences similar to DEBS module 1. Residues are grouped along the axes according to the IC in which they occur, allowing the reader to see the cross-coupling between residues intra-IC and inter-ICs, but meaning that they are not in sequence order. The yellow blocks along the diagonal clearly show high intra-IC coupling, compared to inter-IC coupling, but there is clear evidence of inter-IC coupling, most notably between ICs 13, 5, 11, 2, 3 and 7 in the bottom right corner of the plot. Details of clustering analysis are explained in Methods. Some of the axes labels sit out of line with the other labels due to the very few members in that IC leading to too little space for the label. The colour key represents the scores from the SCA reconstructed without the first eigenvalue and eigenvector.

The whole KR domain is defined by ICs 17, 10 and 21, which form a cluster of ICs that are more cross-coupled to each other than to other ICs (Fig 3) but a catalytic subdomain is defined by IC_3 and to a lesser degree by IC_16. The KR polypeptide is known to fold into two domains, both with short-chain dehydrogenase/reductase folds. One domain is a non-catalytic “structural” domain (KRs), the other is an NADPH binding catalytic domain (KRc) [38]. Whilst the KRc consists of strong inter-residue coevolutionary couplings that result in IC_3, the strong inter-residue couplings of the KRs domain always couple with residues in the KRc (ICs 10, 17, 21), defining the whole KR domain. As well as residues that couple intra-KRc, IC_16 includes couplings with residues from the ACP and the upstream and downstream KS domains, although all the couplings in IC_16 are weak compared to those intra ICs 17, 10, 21 and 3. However, in the cluster analysis, IC_3 clusters with ICs 2 and 5, which consist of residues from the upstream and downstream KS and the ACP, but not with residues in the KRs domain, thus somewhat mimicking what is seen in IC_16.

We were concerned that these results may be an artefact of some sequences in the MSA not having a KR domain but highly similar domain boundaries were detected when only sequences with a domain composition that includes a KR were analysed (S10 Fig IC_10). In that analysis KRc also still comes out as defined by one IC (labelled IC_5, S10 Fig). Interestingly the KRs domain is defined as a separate entity in one IC (IC_20 S10 Fig), but there are only 33 residues from a KRs subdomain of 219 residues, and their coupling scores are low (S10 Fig, lower panel).

It is not possible to tell if these differences are due to a change in the number of sequences analysed or are an artefact arising from the presence/absence of the KR domain in some sequences, but the results of the two analyses are mostly very similar (S1 Table). IC_3 of the full MSA is highly similar to IC_5 from the MSA with a KR in every sequence (S1 Table). Similarly ICs 10, 17 and 21 of the full MSA, which define the KR domain boundaries, were highly similar to ICs 10, 29 and 20, respectively, of the shorter MSA. The analysis of the MSA with a KR in all sequences indicates boundaries for the KR to be R886 to A1356 and for the KRc domain to be P1107 to L1320 (IC 5 of that analysis, S10 Fig)). Considering the analysis of the full MSA, then ICs 10, 17 and 21 of that analysis (Fig 3) together define a boundary for the whole KR, based on highly co-evolved residues, of L896 to R1357 and IC3 defines a boundary to the KRc of G1109 to L1320 compared to the boundaries in the literature [38] of V890 to A1360, KRc starting at position T1110. Thus the domain boundaries identified by analysis of both sequence alignments are almost identical to those established in the literature through laborious experimental work.

IC_3, the IC that identifies the boundaries of KRc, consists of residues that are highly conserved, as do IC_2 and IC_5 (S11 Fig), the ICs that cluster with IC_3 (Fig 3). IC_3 includes the catalytic triad of the KR, K1219, S1243 and Y1256 and the NADPH binding site TGGTGxLG (T1114 to G1121) [39]. Keatinge-Clay and Stroud identified that the adenine ring of NADPH stacks with R1141 and forms hydrogen bonds with D1169 and V1170, which are also detected in IC_3 [38]. Additionally, they showed that the phosphate group of the adenine ribose forms a salt bridge with R1141 and a hydrogen bond with S1142, which are detected in IC_3 and IC_21, respectively. An alpha helix and a loop at its N-terminus, which are close to the active site and referred to as the lid region, are thought to act together with the LDD motif of B-type KR domains, or the conserved tryptophan of A-type KR domains, to determine the stereochemistry of the KR products [9]. The highly conserved second aspartic acid in the LDD motif (D1201) and the residues W1282, T1284, W1285 and G1303 of the lid region are detected in IC_3; whereas the LD of the LDD motif, conserved tryptophan and the rest of lid residues are detected in other ICs as discussed below.

The ICs coupled to IC_3 (ICs 2, 5, 7, 11 and 13) bear highly conserved residues, as well (S12 Fig). IC_2 contains the GXDS motif that is highly conserved in ACPs. The residues of the catalytic triad of the KS1 domain (C173, H308 and H346) are also detected in IC_2. For the KS2 domain while the catalytic residues C1644 and H1819 are detected in IC_2, H1779 is detected in IC_13, which contains highly conserved residues, only from KS2. One intrigue here is how such highly conserved residues can have a coevolutionary signal, however there are sequences where these residues are not conserved, as can be seen in S13 Fig.

The AT together with the linker region immediately C-terminal to it (PAL) seems to be an independently evolving unit. The analysis of the full MSA has the AT consisting of three ICs, 8, 4 and 6, which have residues that cross-couple with each other, as shown by the hierarchical clustering. There is negligible coupling of the residues in these three ICs with residues from any other domains, except some residues from PAL1, which are part of IC_4. (Fig 3). Although ICs 8 and 6 are coupled to residues from the ACP and IC_6 also couples with residues from the KR structural domain, these couplings are very weak compared to those between residues within the AT domain, with e.g. only five residues from KRs and one from KRc being strong enough to pass the statistical threshold for inclusion in IC_6 (Fig 3). IC_4 contains no residues from outside the AT_PAL1 boundary. IC_22, consisting of residues from the post AT linker elements PAL1 and PAL2c, which are very highly coupled, also clusters with ICs 8, 4 and 6 in the robustly selected ICs (Fig 3, lower panel). Although clustering analysis of the full set of ICs places IC_22 closest to IC_34 (S7 Fig), clustered next to ICs 4, 6 and 8, IC_34 was only found in the analysis of the full sequence alignment, i.e. in none of the subsampled analysis, and only has eight residues that fall within our selection criteria for an IC. Thus, it seems that many residues of PAL1 are tightly coupled to the AT domain via IC_4, with residues from PAL1 and PAL2c forming another highly covarying unit that seems to be cross-coupled with the ICs of the AT.

*Trans*-AT sequences were removed from the MSA to test if their lacking an AT influenced the analysis. The alternative alignment led to almost identical boundaries of coevolving residue clusters (S15 Fig). In this *cis*-AT-only analysis, the AT domain is now represented by two ICs. One, *cis*-AT_IC_2,consists of AT and PAL1 with a very few residues from other domains, these latter positions having extremely low coupling to the IC as compared to the AT and PAL1 couplings (S15 Fig). The second IC that has a dominant signal from the AT domain is *cis*-AT_IC_5. Similar to *cis*-AT_IC_2, there are couplings to positions outside of the AT domain but they are weak (S15 FigC). Parenthetically, *cis*-AT_IC_11 has many residues from the AT domain that are moderately coupled, but nonetheless very much more strongly coupled than from residues in other domains, but these also are mostly too weak to pass the test to be included in the IC. All the residues in IC_4 (from the full MSA) are found in *cis*-AT_IC_2, and 78% of the residues from IC_8 are also found in *cis*-AT_IC_2 (S2 Table), suggesting that: (i) the residues in IC_4 are robust to sequence selection, and to some degree also those in IC_8; (ii) the existence of IC_4 and IC_8 as separate ICs is possibly an artefact of the sequence alignment, arising from some mis-alignment of some *trans*-AT sequences. IC_6 and *cis*-AT_IC_5 are also highly similar.

Analysis of the *cis*-AT-only alignment also finds a group of PAL1 residues that form an IC, *cis*-AT_IC_12 together with a few residues from the KAL (S15 FigA). The residues from the KAL are weakly coupled to this IC, but pass the threshold for inclusion in the IC, whereas there are many residues in the AT domain that couple to the IC in the raw analysis (S15 FigB) but don’t pass the threshold for inclusion in the IC. This coupling to *cis*-AT_IC_12 is nonetheless much stronger than from residues in any of the other domains. The PAL2c region couples with many residues in the AT, as well as a few residues in the KRs and KR-ACP linker, as part of *cis*-AT_IC_5, but those coupling outside of the AT region seem few and weak (S15 Fig). However, *cis*-AT ICs 12, 5, 2 and 11 cluster together, again suggesting that the AT and PAL1 form a functional unit, with their relation to PAL2c being less clearly defined.

Taking ICs 8, 4 and 6 of the full MSA together, they define an AT_PAL1 unit of residues V527 to S854, whereas *cis*-AT_IC 2 defines the boundaries of the coevolving unit as residues V527 to R863. The residues of *cis*-AT_IC_11 and *cis*-AT_IC_5 are within the boundaries defined by *cis*-AT_IC_2, except for a few residues of *cis*-AT_IC_5 that are either in the KS, KR or PAL2c region but with low IC scores. Recent experimental work has demonstrated the need for the PAL1, but not PAL2, if one is to successfully replace the AT of DEBS module 6 with that of the equivalent residues from EPOS module 4 [19], corresponding to residues Q524 to P865 in DEBS1. This is consistent with our results here. However, they found that including the KAL region as part of the replacement unit significantly improved yields via improved *K_M_*, which is not evident in our ICs, and this KLA AT_PAL1 also gave functional swaps in other systems (transferrability of the AT_PAL1 unit was only tested in one construct).

Many ICs indicate that the KAL has many weak couplings with the co-joined KS domain, notably in IC_5, suggesting that it is somehow a unit with the KS. However, looking at the scores for IC_26 and IC_1 (S7 Fig), there are also strong intra KAL couplings. IC_1 clusters with ICs 18 and 27, which are defined by strong intra-KS2 and KS1 coupling, respectively. IC_26 clusters with IC_30, which consists of weak residue couplings throughout the sequence but no significant signal in the AT and KRs. IC_26 also clusters with IC_23, but IC_23 was not considered robust. A qualitatively similar picture is seen in the analysis with only *cis-*AT sequences (S15 Fig). It is not clear how to reconcile the coupling of KAL and KS residues with the observation that including the KAL with the AT and PAL1 in AT swap experiments greatly enhances product yield compared to AT or AT_PAL1 constructs, whereas KAL_AT alone has much lower yield than AT_PAL1 [19].

The KS domains seem to be well defined by multiple ICs, with no strong coupling with the ACP-KS linker nor with the AT domain. IC27 has a cluster of strong intra-domain couplings within KS2, whereas IC_18 has strong intra-domain couplings defining KS1 (Fig 3B). IC_13 curiously has extremely high inter-residue couplings in the C-terminal third of KS2. Cluster analysis places it together with ICs 2, 3, 5 and 29, and like them it is highly conserved (S11 Fig). Although it is not clear to us what this means it does not appear to be an artefact of the sequence selection, since it is seen when sampling different numbers of sequences (S5 Fig) and a similar IC is seen in the analysis of the solely *cis*-AT sequences (*cis*-AT_IC_7 in S15 Fig) and in the sequences that strictly have a KR domain (IC_14 in S10 Fig). Based on coevolving residues described by IC_27, the N-terminal boundary of KS2 is M1483 whereas the C-ter is effectively set by where we chose to truncate the sequence alignment. Additionally, including residues from IC_13 defines the N-terminal boundary as D1475. Due to the couplings between the KS1 and its KAL, its C-terminal boundary is not as clear as for the other domains. IC_18 has strong intra-KS1 coupling that defines the C-terminal boundary for KS1 as residue I422. The N-terminal boundary of KS1 is effectively defined by where the sequence alignment was truncated. The MIBIG database [40]entry BGC0000055 annotates the boundaries of KS1 as residues V3 and P426, and of KS2 as I1478 and P1990, i.e. similar to what we see in the ICs. However, the coupling between KS and KAL makes it difficult to objectively distinguish those two segments and thus makes boundary annotation less clear than for other domains.

### 2.4 ICs separate sequences by domain composition and domain specificity

Do different ICs distinguish certain sequence types? E.g. those representing a specific domain composition, or those with certain substrate specificity. The eigen-decomposition of the co-variance matrix results in a matrix with only diagonal values, which represent the variance along the axes of a new coordinate frame. The axes of this new coordinate frame are the eigenvectors. This allows directions with little variance to be discarded and thus the dimensionality of the data can be reduced, making them easier to interpret and store. The original data can then projected onto the eigenvectors to give their position along the new axes, and clusters of related data points can be separated along the axes. Here, we have the problem of projecting categorical data of a protein amino acid sequence, i.e. an alphabetic string such as ADEKLVD, onto an eigenvector (or independent component). To do this, each amino acid at each position in the sequence alignment needs to be converted to a numerical value, which is achieved by looking at the amino acid’s contribution to the inter-column variance of that column in the MSA. This technique was developed by Rivoire et al ([34], see the supplementary information, equation 19 of that paper). The projection is actually onto independent components, which are the result of rotating the eigenvectors to achieve maximal differentiation of clusters along vectors, but the principle of projecting the data back onto these ICs remains the same as projecting data back onto eigenvectors. This projection is referred to as sequence-position mapping and essentially represents the strength of the contribution of each sequence to the covariances that define the IC.

#### 2.4.1 ICs distinguish sequences with different domain compositions

Although the DEBS sequence has KS1_AT_KR_ACP_KS2 domain composition, not all sequences in the MSA have the same domain composition. Some sequences do not have a KR domain and some sequences are from *trans*-AT systems. Additionally, there are sequences that include additional reducing domains. Although these additional reducing domains are removed from the final alignment, the rest of their sequence is kept and thus traces of these domains are likely to be seen in the co-evolution networks. Within the MSA, the most abundant domain compositions detected are: KS1_AT_KR_DH_ACP_KS2 (with 725 sequences), KS1_AT_KR_ACP_KS2 (673), KS1_AT_KR_DH_ER_ACP_KS2 (225), KS1_KR_ACP_KS2 (161), KS1_KR_KS2 (132), and KS1_KR_DH_ACP_KS2 (126).

Sequence-position mapping analysis based on domain composition classification reveals distinct patterns for ICs 4, 6, 8, and 17 (Fig 5 upper panel, S16 Fig). IC_17 consists of strong intra-KR residue couplings and distinguishes the KS1_AT_KR_ACP_KS2 domain composition from KS1_AT_KR_**DH_ER**_ACP_KS2, and KS1_AT_KR_**DH**_ACP_KS2, which suggests that the addition of the DH and ER have some effect on the KR domain’s amino acid composition. A seemingly more trivial separation is that of *cis*-AT and *trans*-AT systems along IC_4 and IC_6, presumably as a result of these ICs consisting predominantly of intra-AT couplings and the cis-AT systems having an incomplete AT sequence, or indeed incorrectly aligned sequence within the AT region (Fig 5 lower panel). IC_8 has two peaks in the sequence-position mapping. The first one has sequences from all domain compositions while the second peak consists of the sequences only from the *cis*-AT systems. We shall see later that the split in the peak correlates with different substrate specificities and so the apparent separation of some *cis*-AT systems reflects this. It is thus evident that the sequence-position mapping can discriminate different domain compositions, although the more interesting results are in its ability to discriminate different subtypes of domain, as we discuss below.

**Fig 5.**
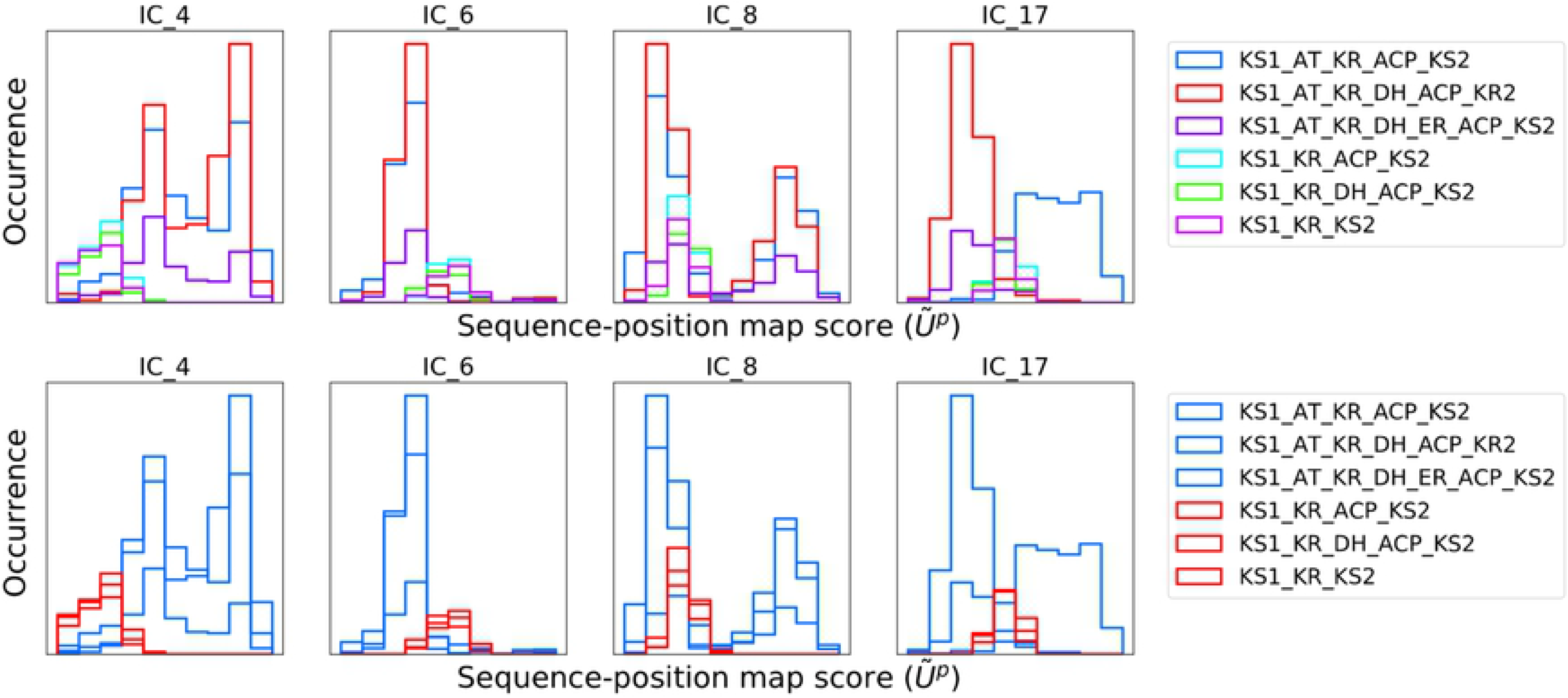
The sequences with different domain compositions revealed different patterns on five sequence maps of ICs. Different domain compositions have diverged amino acids at the corresponding positions in the ICs (upper panel). Conspicuous difference is detected between the *cis-*AT and *trans-*AT systems (lower panel). The horizontal axis represents the projection of each sequence onto the specified independent component, as described in the methods, the vertical axis the number of sequences that have that projection score.

#### 2.4.2 AT domains can be distinguished based on their extender unit specificity

The AT domains in the sequence alignment were classified based on their specificity for their malonyl-CoA, methylmalonyl-CoA, or ethylmalonyl-CoA extender unit, or unclassified, as described in Methods. Sequence-position mapping distinguishes these groups along ICs 1, 4, 6, and 8 (Figs. 6, 8, 10, 11, S17 Fig).

**Fig 6.**
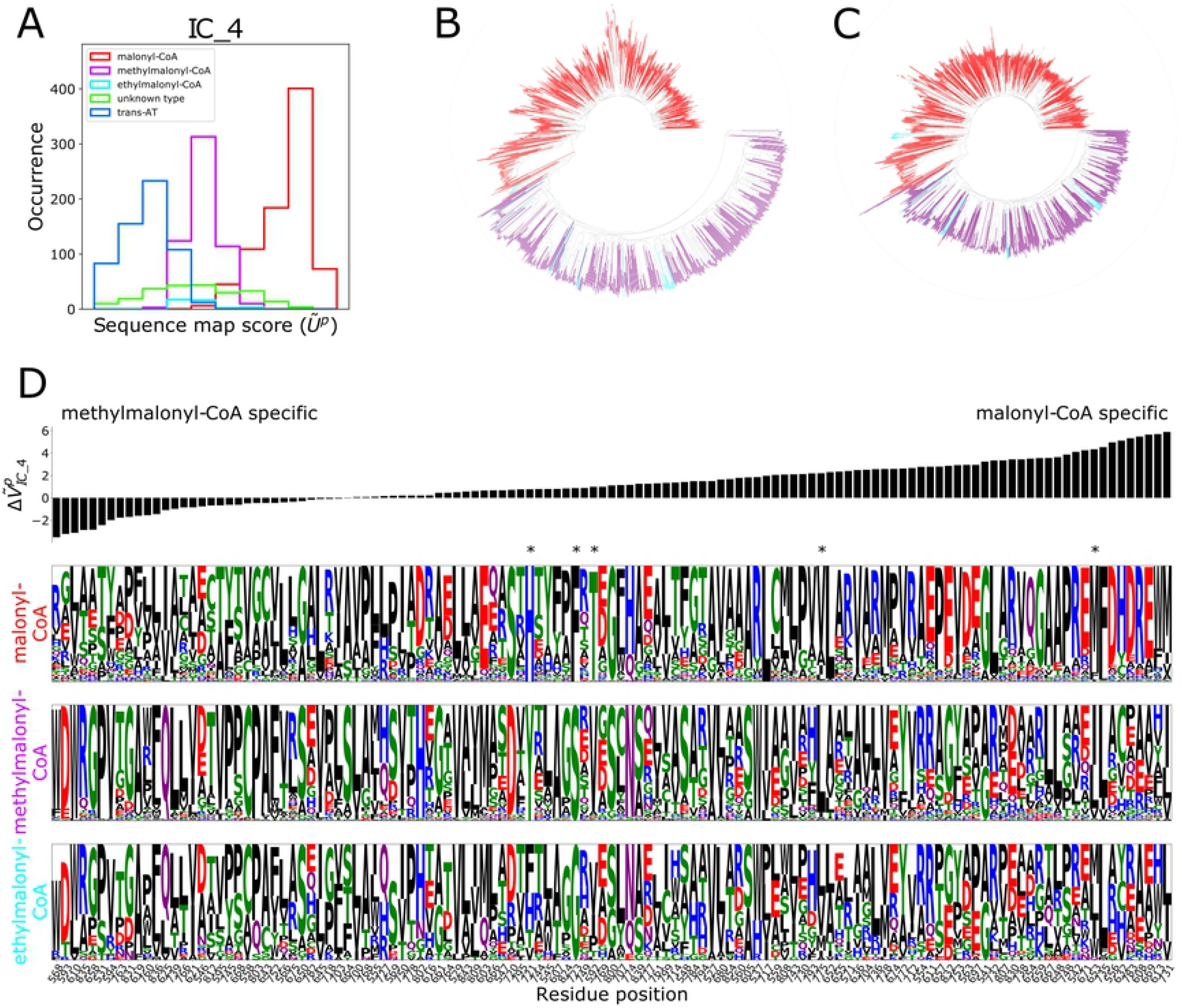
Methylmalonyl-CoA, and malonyl-CoA specific *cis-*AT systems were were distinguished from one another along IC_4. (A) Sequence-position mapping separates the AT domains based on their extender unit specifity. (B) Clustering analysis of the sequences of IC_4 residues distinguishes malonly- and methylmalonyl-specific AT systems supporting the sequence-position mapping patterns of the sequences. The colouring in (B) denotes the same as in (A); *Trans-*AT sequences have been omitted for clarity. (C) Clustering analysis of the whole AT domain reveals a very similar pattern to IC_4. (D) Sequence logos show that sub-types of the AT domains have different amino acid patterns at IC_4 residue positions. The Y and S residues of the YASH motif (methylmalonyl-CoA specificity) and H and F residues of the HAFH motif (malonly-CoA specificity), which provide fingerprints for the two subtypes, are present in this IC and are indicated by * at the top of the column, positions 721 and 723. The * indicate five of the six positions where specificity changing mutations were identified by Zhang et al. [27], as described in the main text. Amino acids are coloured based on their chemical properties. Positions are sorted based on 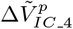 scores, explained in the methods, where 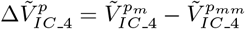, such that the far left of the figure represents residues that we presume to be favourable to methymalonyl selectivity, and those at the far right favourable for malonyl specificity.

Methylmalonyl-CoA and malonyl-CoA specific AT domains show distinct patterns in IC_4 which suggests the residues of this IC should be functional in the extender unit specificity of the AT domain (Fig 6A). The separation that the sequence-position map suggests can also be seen by a dendrogram created by comparing only the residues that contribute to IC_4 (Fig 6B, S18 FigB). From the dendrogram, we can see a clear distinction between malonyl- and methylmalonyl-specific ATs and *trans-*AT sequences. This pattern of the clustering is actually quite similar when we apply sequence clustering on the whole AT domain (boundaries are determined by IC_4 boundaries) (Fig 6C, S18 FigB). That the subtypes separate along IC_4 but not along other ICs suggest that the residues of IC_4 are key indicators of sub-type.

In order to identify the importance of the residues for specific sub-types, we calculated the projection of the sequence covariance of the different subtypes onto the ICs. Conservation weighted covariance matrices for malonly- and methylmalonyl-specific sequences, 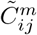 and 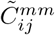 respectively, were projected onto the independent component matrix, calculated from the whole MSA 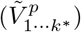.

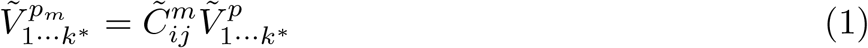

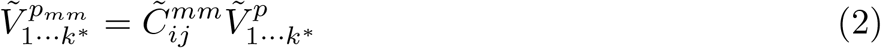

To determine the residues that lead to the malonyl- and methlymalonyl sub-types scoring differently along the different ICs, we calculated the difference between scores of these projections onto the ICs.

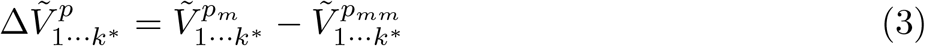

Sorting the positions of IC_4 based on 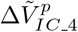 scores gives those that differ most between the two subtypes, with those presumed critical for methylmalonyl specificity at left end of Fig 6D and those for malonyl specificity at the right end. The residues, which are detected in the IC_4 are shown on the structure in Fig 7B, colored based on the 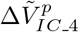 scores changing from green, for positions important for methylmalonyl specificity to blue, positions defining malonyl specificity.

**Fig 7.**
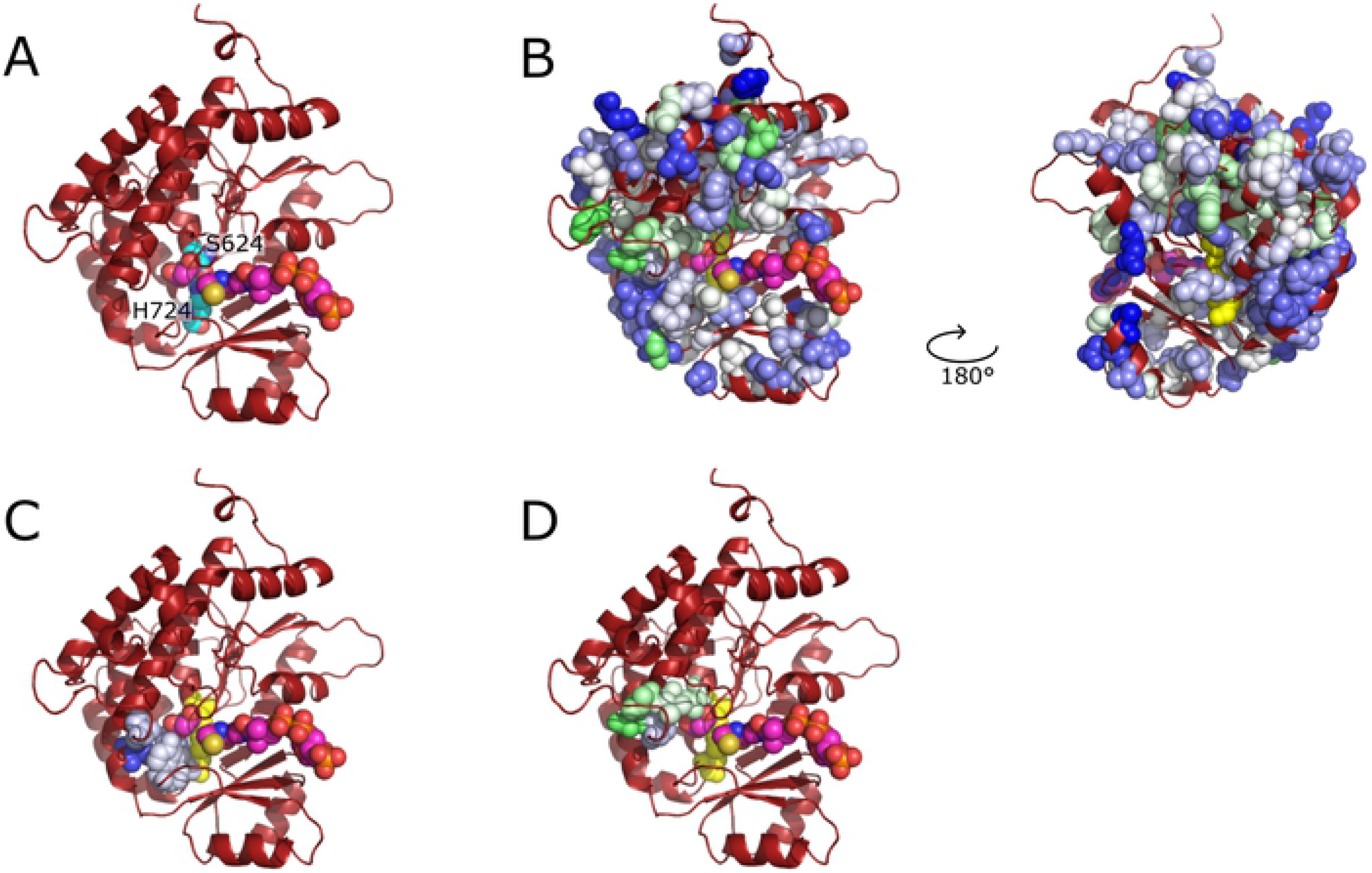
Positions of the residues detected in the IC_4 on the AT domain structure. (A) A model of the structure of AT domain of DEBS module 1 is shown, highlighting the catalytic residues (S624, H724) with sphere respresentations, where cyan represents carbon atoms, red represents oxygen atoms and blue represents nitrogen atoms. Malonyl-CoA is shown to highlight the active site and the access tunnel; the coordinates were modelled based on the substrate position in the crystal structure of FabD (PDB ID: 2g2z [41]), the AT from the fatty acid synthase of *E coli*, without energy optimisation. Carbon atoms of the malonyl-CoA are shown magenta, oxygen atoms are shown red, nitrogen atoms are shown blue, phosphorous atoms are shown orange and the sulphur atom is shown yellow. (B) Residues detected in IC_4 are shown with spheres, which are colored based on the 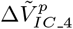 scores. Darker green represents lower 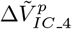 score indicating methlymalonyl specificity, whereas darker blue represents higher 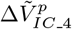 score indicating malonyl specificity. All atoms of the catalytic residues are shown yellow. (C) Residues identified by Zhang et al. [27] and detected in IC_4 are shown as spheres. (D) Additional residues that have been experimentally associated with AT domain specificity and detected in IC_4 are shown as spheres (V592, D593, V594, L595 and Q625 in the text although 595 is valine in the DEBS module 1 sequence used to model this AT).

The sequence logos show that positions that scored highly for a subtype are mostly conserved for that sub-type but not for the other sub-types. There is an exception at position 760 that has a score in favour of methylmalonyl-specificity yet the conservation is not high. And a similar, but more conserved pattern is seen for position 763. In order to investigate these positions further, 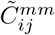 was analysed revealing that residue 760 has the highest coupling with residue 753. The spatial positions of 753, 760 and 763 on the 3D structure are close to each other (S19 FigB). The most abundant pairs for positions 760 and 753 in methylmalonyl-specific sequences are Trp-Phe, Leu-Tyr and Arg-Tyr suggesting that the interaction between these two positions may be important and evolutionarily constrained (S19 FigC); whereas, a similar pattern is not observed for malonyl-specific sequences (S19 FigD). The C_*β*_-C_*β*_ distance between residues 760 and 763 is 5.2 Å.

The Y and S residues of the YASH motif and H and F residues of the HAFH motif, which provide the distinction between the fingerprint motifs, are detected in this IC (positions 721 for Y/H and 723 for S/F, marked with stars in Fig 6). Site-directed mutagenesis studies on the YASH and HAFH motifs, aiming to switch the extender unit specificity, generally result in promiscuous domains which can accept both methylmalonyl-CoA and malonyl-CoA as an extender unit. Further, kinetic analysis shows that the cause of the promiscuity is not based on an increased affinity for a non-native extender unit but rather a decrease in the capability of accepting the native one. A recent study by Zhang et al. performed experiments on salinomycin polyketide synthase, targeting additional residues on malonyl, methylmalonyl and ethylmalonyl specific ATs besides the YASH/HAFH motifs [27]. After performing structural analysis and molecular dynamics simulations, they determined that hydrophobic residues at positions V592, I653, M662 and L775 (DEBS1 module 1 numbering) are critical for substrate specificity. While mutating the residues of the YASH/HASH motifs did not provide a switch from one specificity to another, additional mutations at the four newly identified positions successfully switched between malonyl/methylmalonyl/ethylmalonyl specificity. The SalAT2 H276Y/F278S/I177Q/M205L/V327 mutant showed a change in substrate specificity from malonyl CoA to methylmalonyl CoA, and reverse mutations, i.e. Y288H/S290F/Q189I/L217M/L341V, in SalAT8 changed its specificty from methyl malonyl CoA to malonyl CoA. Consistent with these results, except the residue at 662 (which is in IC_6 and found highly conserved as Met in all specificities, malonyl/methylmalonyl/ethylmalonyl, but Val in the ethylmalonyl specific AT of the salinomycin pathway), the other positions were detected in the IC_4 (Fig 7C). Thus, residues important for specificity, which were identified by analysis of structural dynamics, can be identified directly from the sectors analysis of sequence data.

As well as the YASH/HAFH motifs and residues identified by Zhang et al., IC_4 also encompasses D593, V594, L595, and Q625, which have all been shown to affect the extender unit specificity of ATs. In AT4 of the DEBS pathway, mutating RVDVL to DTLYA allows the AT to accept a malonyl extender unit as well as methylmalony [24, 42]. D593, V594, and L595 are second, seventh and nineteenth from the left in Fig 6D, which thus predicts them as particularly likely to affect specificity, and V592 is very much in the middle of the plot. The V to T mutation is predicted by Fig 6D to weakly favour malonly specificity, T being conserved in malonly specific ATs, whereas DVL to LYA should disfavour methylmalonyl specificity. Q625 is adjacent to the catalytic serine in the GHSXG motif, which was detected as a branched hydrophobic residue (Val or Leu) in malonyl-specific ATs but as methionine or glutamine in other AT types [42, 43]. Q625, is 13 positions from the left side of Fig5D and very highly conserved in methylmalonyl specifc ATs. R591, the first residue of the RVDVL motif, occurs in IC_6, which doesn’t discriminate substrate specificity, as discussed later, and is conserved as either R or D. Thus IC_4, which is determined purely by sequence analysis, identifies five residues that have previously been shown to be important for AT extender unit specificity (Fig 7D) and highlights further residues that could be important.

Although it is not impossible for the method to have detected these key residues by chance, it seems unlikely. For example, in the Zhang et al. [27] study there are six residues that were experimentally shown as important for the extender unit specificity. We found five of them in the same IC (IC_4). As we could detect five of the six residues, the probability of detecting them by chance in an IC with a length of 123 amino acids (length of IC_4) from a domain sequence whose length is 342 (AT domain + PAL1 region) can be calculated via hypergeometric distribution, 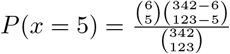, which is equal to 0.022. This low probability indicates finding five of the six experimentally determined functional residues in an IC with 123 residues by chance is quite small.

Historically, it has proved difficult to identify the residues that do provide substrate specificity to the different ATs, indeed identifying residues important for catalytic activity and residue specificity is a general problem in protein design. Directed evolution has often demonstrated that residues far from the active site can have a large effect on these properties and it is not understood why, which makes it difficult to identify such residues. The residues identified here are also seen to be distributed close to and far from the active site cleft (Fig 7), suggesting that this may be a way to identify such residues. Some caution is required, since the residues that have been identified are generally highly conserved within subtypes and the correlations seen might be driven by unimportant changes in common ancestors, i.e. tied to phylogeny but not function. However, support for interpreting the residues as functionally important is provided by the number of residues in IC_4 that have already been identified as important for functionality, as noted above. Thus, the residues identified at the left and right sides of Fig 6D and at the left side of Fig 8C (discussed below) are prime targets for mutagenesis work to change the substrate specificity of AT domains, although the implication of the method here is that the correct network of residue types needs to be incorporated, and not simply isolated mutations [44].

**Fig 8.**
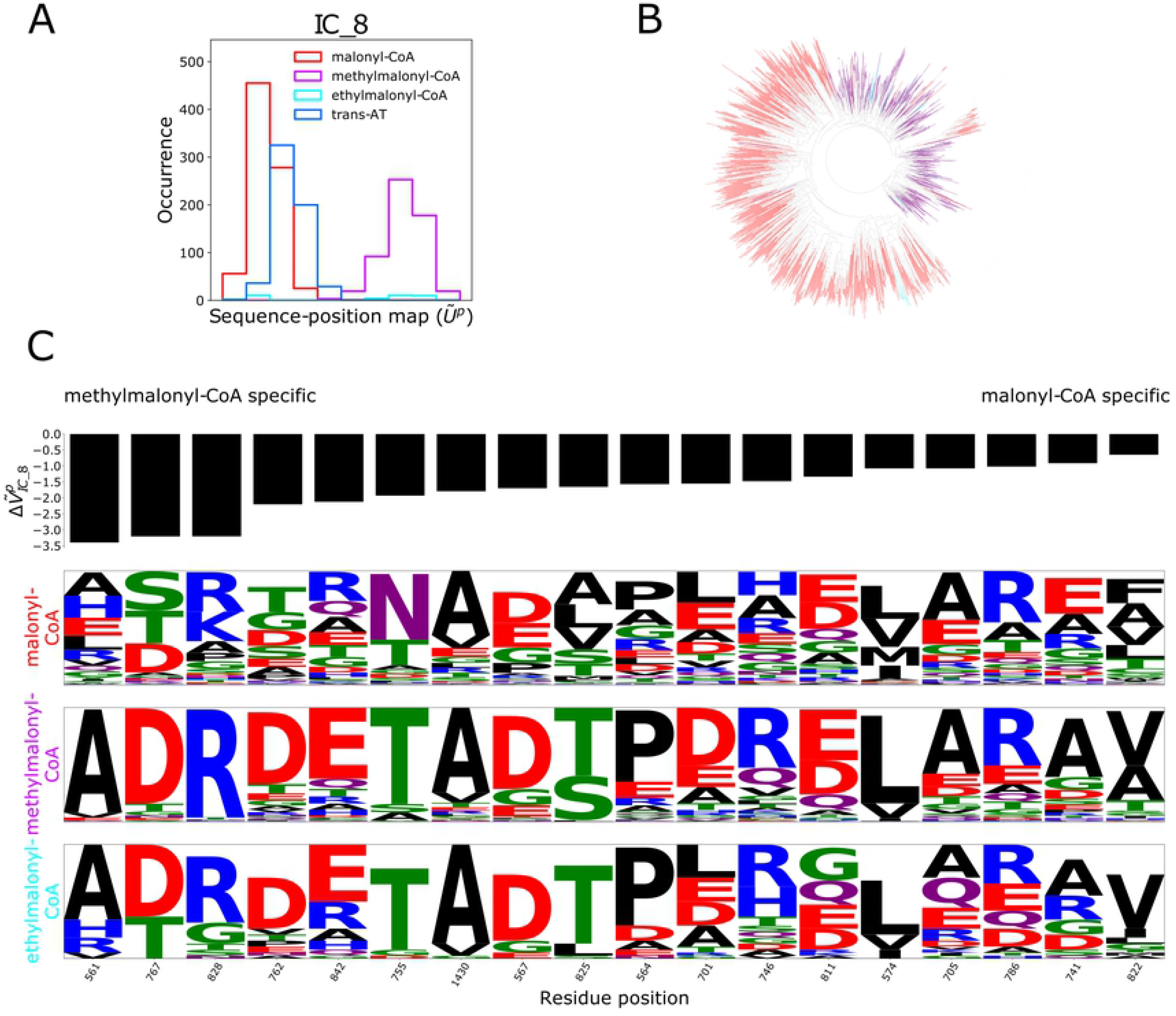
The residues of IC_8 are more conserved in methylmalonyl- and ethylmalonyl- specific ATs compared to malonyl- specific ATs. (A) Sequence-position map of IC_8 separates methylmalonyl- (and ethylmalonyl-) specific ATs from malonyl-specific ATs. (B) Clustering analysis of sequences of IC_8 residues clusters methylmalonyl-specific ATs in the same branch, the colouring scheme is the same as in (A). (C) Sequence logos of IC_8 residues reveal that those positions are more conserved in methylmalonyl- and ethylmalonyl- specific AT domains, contrary to malonyl-CoA specific ATs. Positions are sorted based on the difference between AT sequences with malonyl and methyl malonyl specificity where 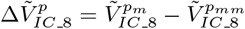.

Sequence-position mapping based on the AT extender unit specificity also clarifies the ambiguity of the double peak in IC_8 arising from the domain composition mapping (Fig 8A). The second peak is composed of only methylmalonyl specific ATs while the first peak bears a mixture of the rest. The distinction between the methylmalonyl- (and ethylmalonyl-) specific ATs and the others is also clear in pairwise comparison of the IC_8 residues’ sequence. (Fig 8B, S18 FigC).

The residues of IC8 are much more highly conserved in the methylmalonyl and ethyl malonyl specific AT domains than within the malonyl specific ATs (Fig 8C). We are not aware of any experimental work to support or disagree with a role for these residues in methylmalonyl specificity, yet the separation of the methylmalonyl specific AT sequences for the malonyl specific ones is much clearer along IC_8 than for any other IC, which is consistent with the large values of the 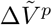 between methylmalonyl and malonyl specific ATs. It seems worth reminding the reader that the residues in IC_8 and IC_6 were present together in *cis*-AT_IC_2, from our analysis solely of *cis*-AT sequences, consistent with their both having a similar role, which seems most likely to be substrate specificity, and they cluster together in the hierarchical clustering, indicating that there is some coupling between residues in the two ICs. The residues of IC_8 are shown on a model structure in Fig 9

**Fig 9.**
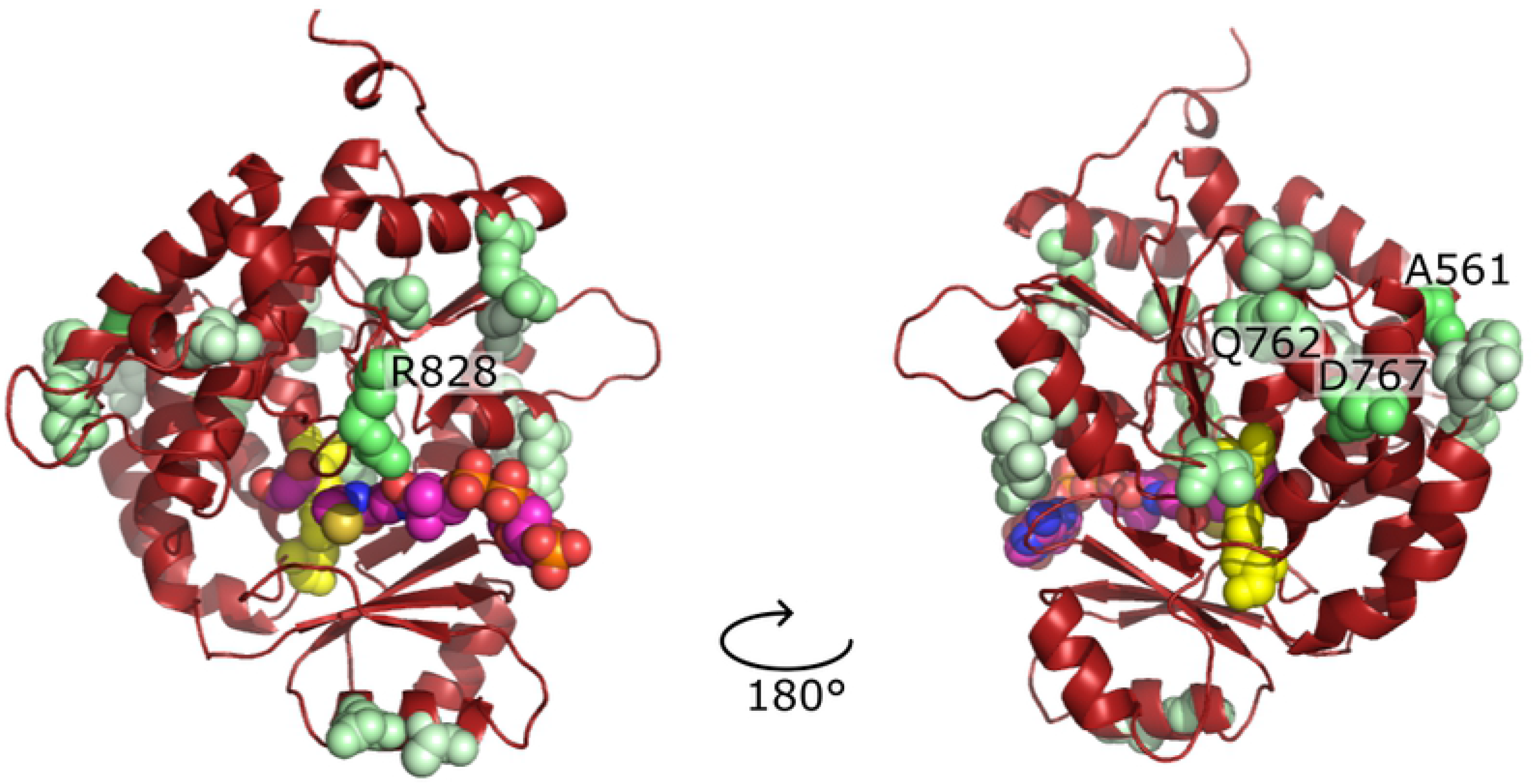
The positions on the structure of the AT domain of the residues of IC_8. The residues of IC_8 are shown as spheres on a model of the AT domain from DEBS module 1. IC_8 residues are colored based on 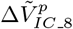 scores where darker green represents the strongest methlymalonyl specificity. Residues with the largest magnitude 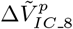 scores (R828 in the front view, and A561, D767 and Q762 in the back view) are labeled. Malonyl-CoA is shown to highlight the active site and the access tunnel, as described in Fig 7. Carbon atoms of the malonyl-CoA are shown magenta, oxygen atoms are shown red, nitrogen atoms are shown blue, phosphorous atoms are shown orange and the sulphur atom is shown yellow. All atoms of the catalytic residues are shown yellow.

Highly conserved residues in both malonyl-CoA and methylmalonyl-CoA specific subtypes are detected in the IC_6 (Fig 10B). Catalytic residues (S624, H724) are also detected in this IC. This suggests that the residues detected in IC_6 are critical for the proper function of the AT domain irrespective of the extender unit specificity.

**Fig 10.**
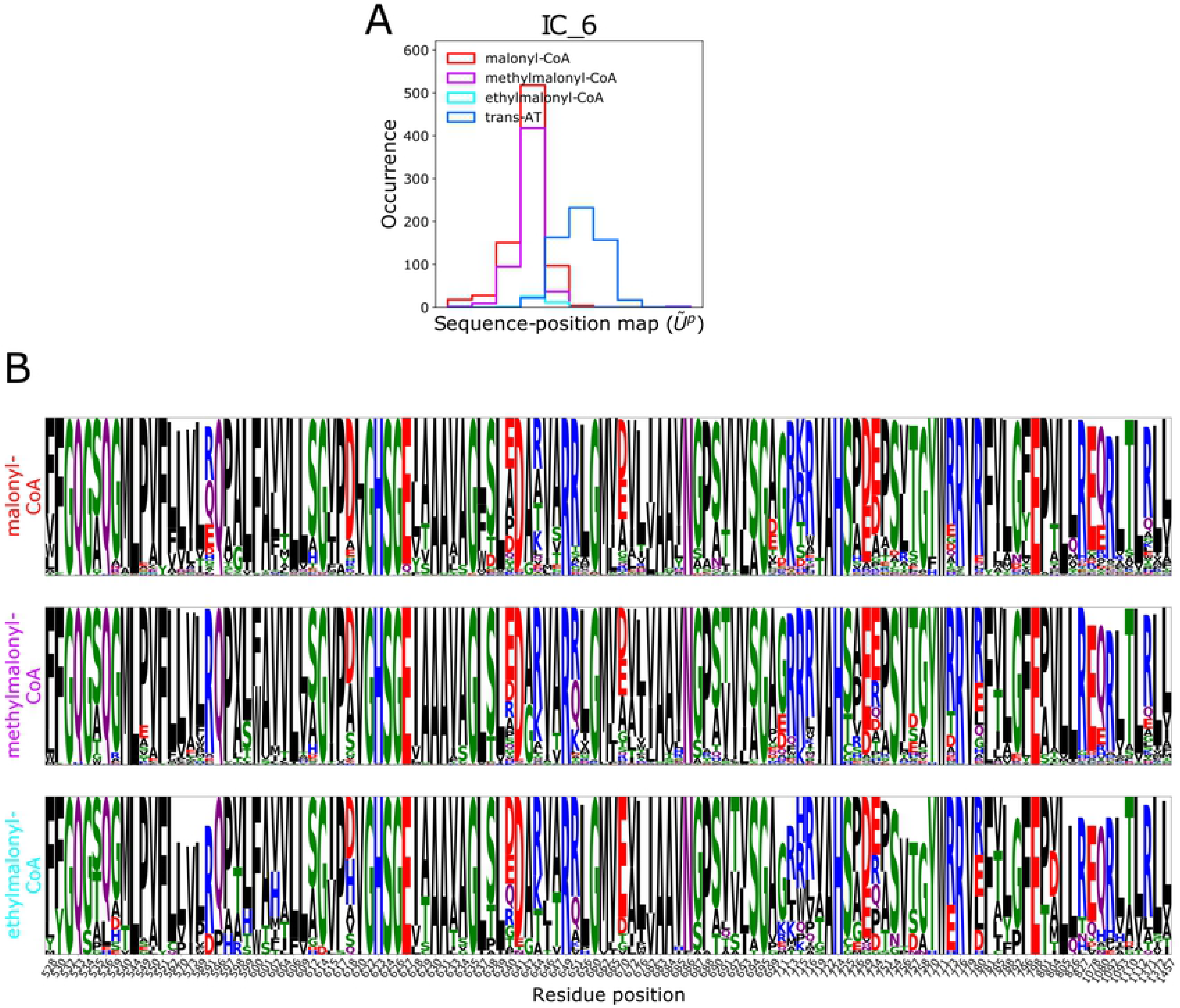
IC_6 consists of Highly conserved positions of the AT domain, including the catalytic residues. (A) Sequence-position mapping with IC_6 doesn’t separate *cis-*AT sequences by substrate subtype. (B) Sequence logos show the high conservation of the residues in IC_6, including catalytic residues (S624, H724), across methylmalonyl-CoA, malonyl-CoA and ethylmalonyl-CoA specific AT domains.

#### 2.4.3 Residues that distinguish *trans*-AT and *cis*-AT systems

A large number, but not all, *trans*-AT sequences separate from the other sequences by sequence position mapping along IC_1 (Fig 11A), which consists of residues from both of the KS domains, KAL, ACP and KRc, but not the KRs, ACP linkers, AT or the PAL1. These residues show very similar variation across the *cis*-AT sequences, but see different residues at these positions in the *trans*-AT sequences, and slightly higher conservation (Fig 11C).

**Fig 11.**
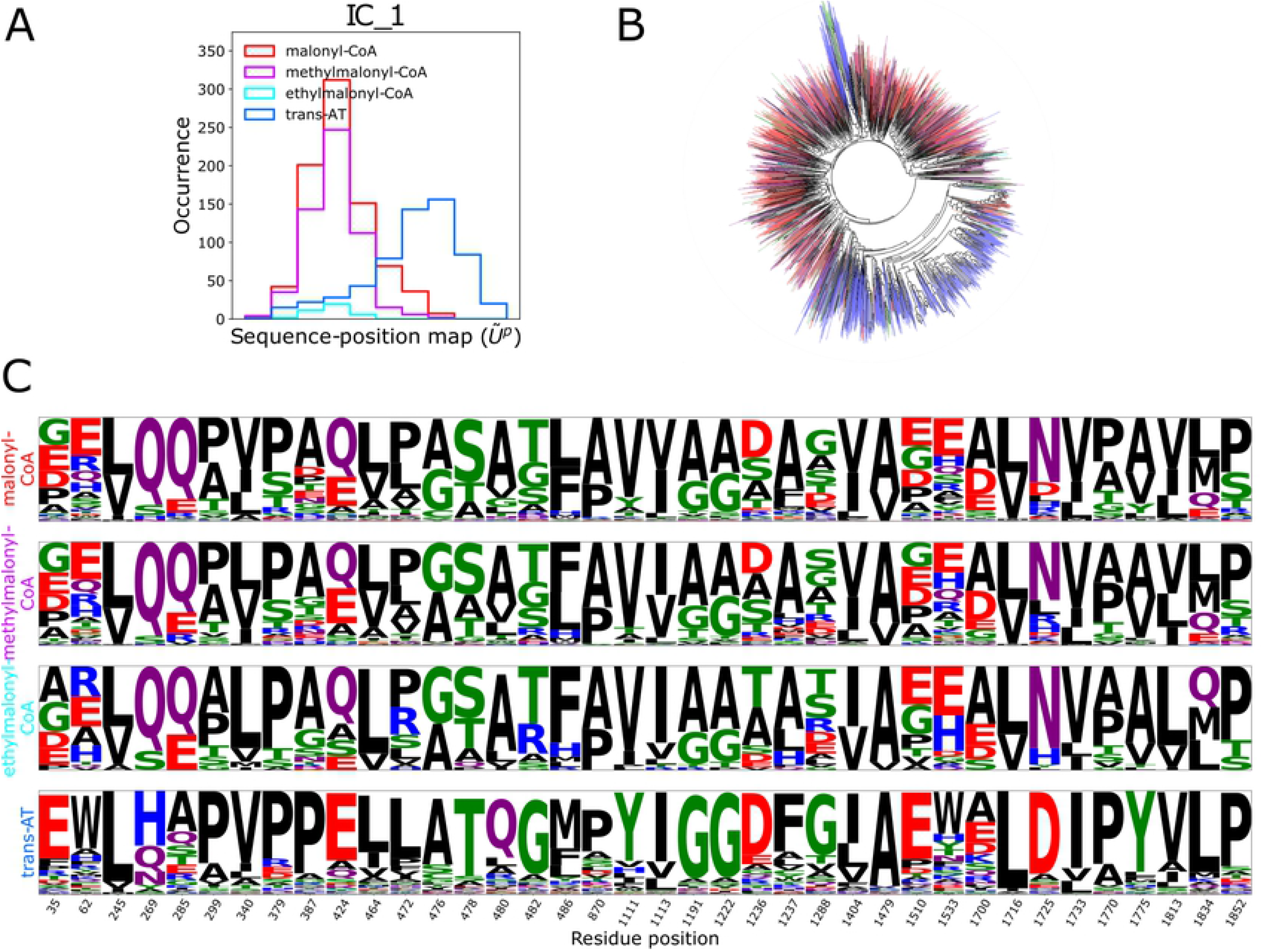
Different sequence patterns outside of the AT domain in *cis-* and *trans-* AT systems are detected in IC_1. (A) Sequence-position map of IC_1 separates *cis-* and *trans-* AT systems. (B) Clustering analysis of sequences of IC_1 residues clusters *trans-*AT sequences, the colour scheme is the same as (A). (C) Sequence logos shows the residues of IC_1 having different amino acid types in *trans-*AT and *cis-*AT seqeuences.

#### 2.4.4 KR domain types can be distinguished by the independent components

For most ICs the different KR types show a very similar distribution, which is to be expected since most ICs do not encapsulate the covariance of residues within the KR, but along some ICs the distributions skew such that the KR types separate into A, B and C types. The distribution of C type KR domains in IC_3 are skewed to the left compared to other KR types, as it also is to some degree in IC_10, and the distribution of C type domains along IC_21 is slightly skewed to the right compared to other KR types (S20 Fig). The residues covarying in IC_3 are from the KRc and KRs subdomains and include the conserved catalytic triad, so it is congruent with this that the non-reducing C type KR domains should have a different coevolutionary profile for this set of residues compared to the reducing KR types. Similarly IC_10 and IC_21 consist of covariance from residues in KRc and the KRs. In the hierarchical clustering ICs 3, 10 and 21 are closely related (Fig 3). In contrast IC_16, which also contains coupled residues from the KRc, albeit more weakly coupled than in IC_3, does not show a skew in the distribution of C type sequences compared to the other KR types.

Thus C type residues can be separated from other KR types by sectors analyis. This might be anticipated since we need only detect the non-functional substitution of the residues required for the reduction of the *β*-keto group. KR domains of the C2 type do need to maintain the capability to bind the substrate and to epimerise the alpha carbon. We did not attempt to separate the C1 and C2 types along the ICs, since we are not aware of any existing sequence signature that we could use to characterise the two types of sequence.

The results suggest that A and B type KR domains can also be separated along the ICs, although there are too few examples of B2 type for any conclusions to be drawn for these. IC_17 shows a skew to the right for A type and to the left for B1 type (Fig 12). C type have no skew along this IC. A similar although weaker skew is seen when the sequences are projected onto IC_10, which as noted above also shows a left hand skew for C type KR domains (S20 Fig). IC_10 and 17 are next to each other in the hierarchical cluster analysis and are two of the most closely coupled ICs (Fig 3). This suggests that IC_17 and, to a lesser extent, IC_10 specify the residues that determine whether the beta-hydroxyl is L or D configured.

**Fig 12.**
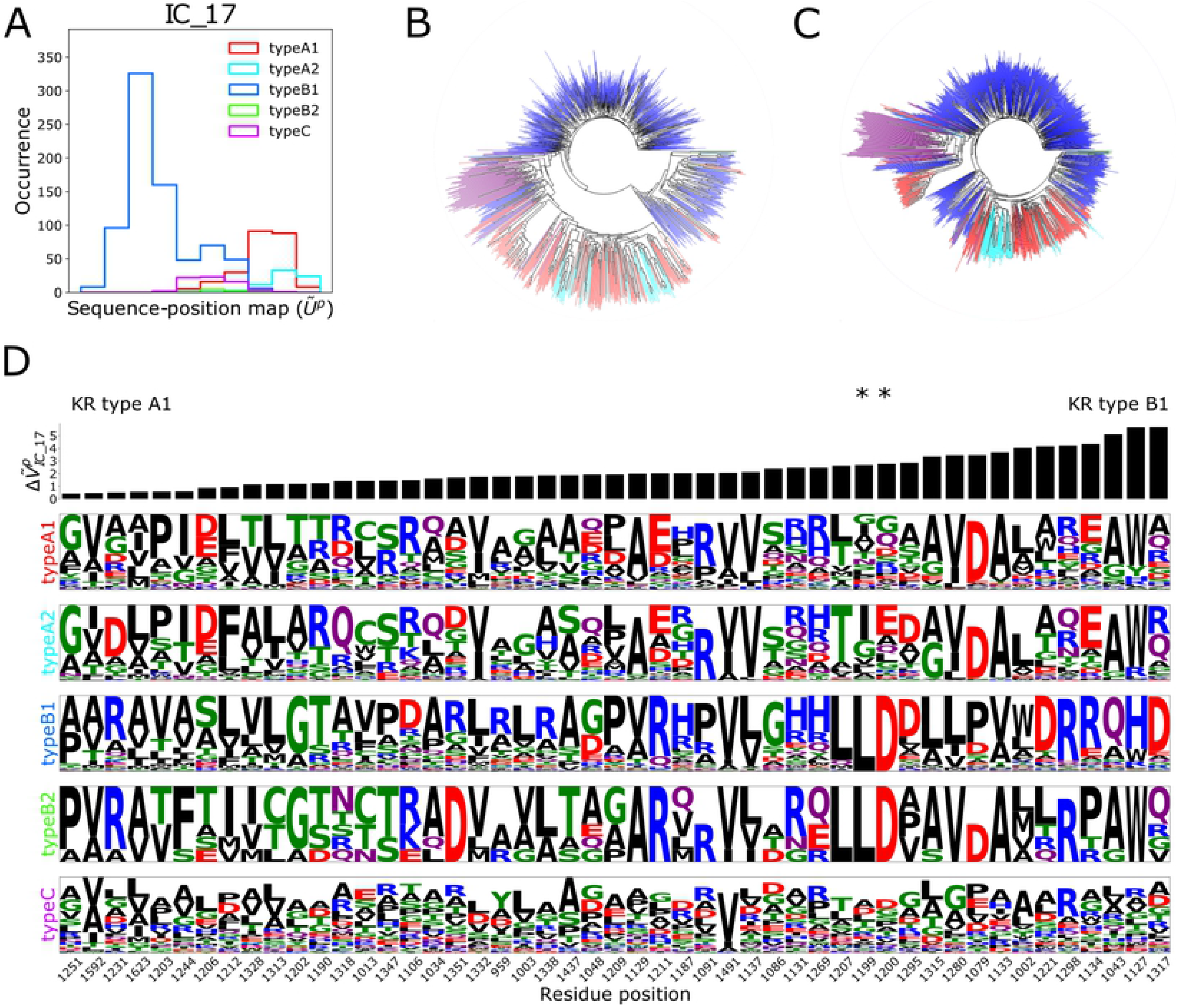
Different sequence patterns for type-B and type-A KR domains are detected in IC_17. (A) The sequence-position map separates the KR domains based on their sub-types. (B) Clustering analysis of the sequences of IC_17 residues distinguishes type A and type B KR domains supporting the sequence-position mapping patterns of the sequences. (C) Clustering analysis of the whole KR domain reveals a similar clustering pattern to that based solely on the residues of IC_17. (D) Sequence logos show that positions detected in IC_17 are more conserved in type B ketoreductases compared to type A. Positions are sorted based on 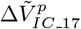 scores. Residues L1199 and D1200 of the LDD motif, which is a finger print for type B KR domains, occur towards the right of the figure. W1248, the fingerprint residue for A type KR domains, is not present, being found in IC_10.

B type KR domains have a sequence fingerprint of an LDD motif, of which KR1 of the DEBS pathway, the one we use in our search query here, is an example. The LD of this motif is present in IC_17, however the D following the L is not well conserved and is not thought to be particularly important. The final D of the motif has been shown to be important and is hypothesised to control the direction of entry of the substrate into the active site, which is thought to control the direction from which the hydride attacks the carbonyl group and thus the chirality of the resulting beta-hydroxyl [38]. The mutant D1201A (according to the numbering scheme we use here, the last D of the LDD motif) reduced the production of the natural product by 40%, and D1201A/F1244G reduced production to 10%, also implicating F1244 as an important residue [38]. D1201 is detected in IC_3, which contains the highly conserved positions of KRc (S11 Fig) and F1244 is detected in IC_17.

Type A KR domains have no LDD domain but rather have a conserved tryptophan and these two changes, as compared to B type, are thought to lead to the substrate entering with the opposite face of its beta-carbonyl facing the NADPH, and thus to the L stereo-configuration of the resulting beta-hydroxyl group. This key tryptophan, located at position 1248, is in IC_10, which also includes residues from a loop-helix lid region that closes the top of the active site cavity after substrate binding. Residues A1286, G1287, A1291, from the loop, and F1299, R1300 and H1302, from the last turn of the helix, are in IC_10. The helix of this region is thought to be also involved in directing the substrate into the active site, with residues V1295 and R1298 (inferred from the structure of first ketoreductase of the tylosin PKS, pdb ID 2Z5L) [9], which are from two consecutive turns of the helix and most likely point from the helix into the space where the substrate is thought pass in A type KR domains [9], being in IC_17. Key conserved residues at the N-terminus of the loop and C-terminus of the helix are in IC_3 (A1281, T1284, W1285 and G1303). For completeness, G1283 is in IC_16, S1288 is in IC_1 and G1289 and M1290 are in IC_28.

Sequence position mapping of IC_17 distinguishes A1/A2 type KR domains from B1/B2 type KR domains (Fig 12A). The similarity between the sequences when considering only those residues contributing to IC_17 is shown in Fig 12C as a dendrogram. Although the clustering pattern of IC_17 residues resembles the dendrogram from clustering whole KR domain sequences (Fig 12B) differences in the distribution of different KR types can be seen.

Similar to our analysis of AT domains, we identified the residues of IC_17 that differentiate A1 from B1 ketoreductases, by calculating the difference of the scores of the sequence subtypes projected onto the IC. Since the number of sequences of type B2 and type A2 are low, we applied this approach only on types B1 and A1.

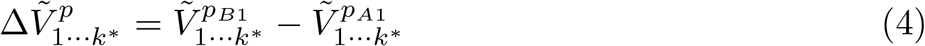

The residues in the IC were sorted according to the 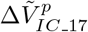 scores (Fig 12D). The LD residues of the LDD motif are close to the right end of the plot supporting the importance of these residues for type B specification. Interestingly, we detect 12 residues with higher scores than L1199 and D1200 residues, suggesting these 12 positions should be taken into consideration for experimental studies to switch the KR type to B2.

### 2.5 Effects of the method of alignment on results

One methodological challenge is to determine which sequences to keep in the alignment, as different alignments may lead to different IC sets. Although differences in the alignments used here resulted in variations in the ICs, the interpretation of the results is not qualitatively changed (Figs. S10 Fig, S15 Fig, S1 Table, S2 Table). Additionally, there are some unexpected residues in certain regions of the MSA, possibly causing some noise in the results. For example, there are some residues in the AT region of the sequences labeled as *trans-*AT while no residue would be expected to be detected in that region. These residues in the *trans*-AT sequences are thought to arise either from the presence of remnants of the AT domain or parts of other domains that were misaligned. In order to investigate how the results would change if the MSA were generated in an alternative way, sequences were grouped according to their domain composition and aligned only with sequences in their group. Groups with fewer than ten sequences were discarded. The groups of aligned sequences were then aligned with respect to each other. ICs calculated from the MSA generated by this domain composition grouping protocol(S21 Fig) were compared to the ICs determined from the full MSA (S3 Table). Although the overall similarities between the ICs are not very high, the ICs that define the domain boundary and show sequence divergence in different domain sub-types have high similarity with the ICs of the MSA generated based on domain composition (S4 Table), suggesting that even though the MSA varies, ICs that carry important information can be detected.

## 3 Conclusion

Key targets for engineering *cis*-AT systems such as the DEBS pathway are the loading module and the AT and KR domains. Substitution or modification of the loading domain can lead to a change of the starting building block. Changing the substrate specificity of an AT domain can change the extender unit, e.g. changing specificity from malonyl to methyl malonyl will add a methyl group to the alpha position. Changing KR domains can potentially alter all and any of the stereo centres in the backbone of the polyketide. During each elongation step a KR can determine the configuration of the stereo centre at the beta position, where there is a hydroxyl, and at the alpha position, via their epimerisation capability. Since the next elongation step leads to the carbon that was previously in the alpha position becoming the gamma position and the previous beta position becoming the delta, then the KR domains have the capability of determining the configuration of every stereo centre on the backbone. Engineering these parts of a PKS can thus have a massive impact on the polyketide that can be synthesised.

With SCA/sectors analysis we can detect functional domain boundaries consistent with experimental studies as well as residues strongly associated with subtype specificity. This gives one explanation for the importance of domain boundary optimisation for the successful swaps [19], since residues within the domain boundaries have clearly been under selective pressure to function together. Furthermore, sequence-position mapping analysis identified groups of residues that distinguish subtypes of domains, with the analysis showing the inclusion of experimentally validated residues but also implicating further residues as important. This is consistent with experimental results since mutating only a few residues has not yet switched a domain from one subtype to another.

The challenge, as regards re-engineering for novel purpose a multidomain protein such as a polyketide synthase, is not to identify the functional domains; hidden Markov models, BLAST and similar sequence searching tools can perform this job well. The challenge is to know exactly where the functional boundaries of these domains lie, so that domains can be spliced from one system into another to generate novel functionality. In the case of polyketide synthases the KR domains were originally only annoted for the KRc segment and the structural subdomain of the KR was completely ignored [38], yet the coevolutionary analysis here marks the KRs and KRc as one clearly co-evolving unit very much distinct from the rest of the module. Similarly, the analysis also highlights the AT_PAL as a distinct coevolving unit, which has been shown in experiment to be critical for swapping an exogenous AT for the native AT of a module. The coupling we see here between the residues in the C-ter of KS2 is intriguing and it is unclear what it represents functionally, but it may provide some insight into how to reengineer KS domains.

However, some parts of the analysis point to continued challenges to re-engineering, even if domain boundaries and active site specificities can be well defined. IC_23, IC_26 and IC_30 show weak coupling across all domains and cross-couple with each other (S7 Fig), although ICs 23 and 30 are not consistent across bootstrap calculations. Nonetheless, this points towards weak levels of co-variance across all domains in the module, suggesting that optimal function of a re-engineered PKS may always require some level of directed evolution.

Here we have highlighted KR and AT boundaries and residues associated with subtype specificity but an attentive reader may spot further associations in the data. For example, IC_8 and IC_4 are determined by the covariance of residues in the AT domain, but show separation of A2 type KR sequences. IC_8 is bimodal, with the left hand peak correlating with malonyl extender units and the right hand peak correlating with the ATs activating methyl or ethyl malonyl extender units. The distribution of A2 type KR sequences along IC8 disproportionately associate with the methyl malonyl and ethyl malonyl AT types. Similarly, projecting the sequences onto IC4 shows a leftward skew in the distribution of A2 type KR sequences, which again correlates with methyl malonyl and ethyl malonyl AT types. This assocation is presumably because there is no need to epimerise the alpha carbon if it has two hydrogen atoms attached, as is the case for malonyl extender units. Sectors analysis has been found to provide some insight into allosteric regulation [45] and future work looking at sectors in PKSs may give clues as to how they coordinate the biosynthetic process across multiple domains and modules.

Beyond the interpretation of results in PKS systems, and the potential to break a bottle neck in the research in that field by identifying residues that correlate with subtype specificity, we have also demonstrated the benefits of using bootstrapping to test the robustness of results, which is likely to be important in the application of these methods to other multi-domain protein families. There seems to be no obvious method for *a priori* determining the number of sequences needed for a reliable sectors analysis, but bootstrapping provides a way of having confidence in the output. We have also shown how to adapt the existing method to predict the likely importance of different residues in their contributions to subtype specificity.

The SCA/Sectors analysis applied here is an unsupervised method for discovering structure within protein sequences, as are related techniques such as DCA [46, 47], and may benefit from coupling with supervised deep learning techniques or the recent development of unsupervised pre-training methods arising in the field of deep learning [48], or through other machine learning techniques. Despite the success of deep learning techniques during the last 15 years, there has been increasing interest in unsupervised machine learning techniques as a potential means to improve the performance of the supervised learning that typically constitutes deep learning. In the structure/sequence analysis of proteins, such unsupervised methods have been used as an input for supervised deep learning to predict inter-residue distances and contacts [49–52], but more recently such models have been incorporated directly into deep networks [53, 54], allowing a generalised learning across protein families and thus improved predictions where protein sequences are few in number. Since the depth of the sequence alignment was clearly important in the analysis presented in this paper, the incorporation of such techniques into neural networks proposes exciting possibilities for the future. Beyond this other methods for calculating groups of co-evolving residues, not investigated here, already exist [55, 56] and using unsupervised training of deep networks also seems to reveal information about the structure of proteins [57]. However, the aforementioned methods have yet to be tested against experiment in the way that SCA/sectors analysis has. A particular interest would be whether they can recapitulate the success of SCA/sectors in predicting pathways of allosteric connectivity through the protein, which is of interest not only for studying protein function and evolution, but also potentially for drug discovery [58]. The results presented here, for the PKS system, support the SCA/Sectors analysis as already able to provide predictions of domain boundaries and important functional residues.

## 4 Methods

### 4.1 Generating the Multiple Sequence Alignment

The sequence of DEBS1 module 1 together with the KS of module 2 was selected as the query sequence (UniprotID:Q03131) (S1 Fig). Homologous sequences were detected with HHblits [59], from which a multiple sequence alignment was generated. For each sequence in the alignment, pfam domains [60] were determined via hmmscan [61] with the e-value set to 0.001. Sequences whose original module includes domains other than KS, AT, KR, DH, ER, TE and ACP were removed from the alignment. After preprocessing the MSA to improve its quality, as described in ref. [34], e.g. to remove highly gapped columns and sequence fragments, the final alignment included 2303 sequences. From the processed alignment, only the sequences with at least two KS domains and an ACP domain were kept in the final alignment, ending up with 2245 sequences.

The MSA was preprocessed with the scaProcessMSA.py script from the SCA analysis tool [34] and the reference sequence was set to the DEBS1 query sequence by–refindex 0 command. The second and third steps of the analysis, which apply the SCA method and determine the independent components, were performed with the scaCore.py and scaSector.py scripts [34].

### 4.2 AT Domain Classification

Sequences that did not give a hit for the AT domain with hmmscan were labelled as *trans-*AT systems; whereas, the ones that gave a hit for the AT domain were labelled as *cis-*AT sequences. The *cis-*AT sequences in the alignment were classified as specific for malonyl-CoA, methylmalonyl-CoA and ethylmalonyl-CoA based on their fingerprint motifs HAFH, YASH and (T/F/V/H)AGH, respectively [43]. AT sequences not bearing any of these motifs were labelled as ‘Unclassified’.

### 4.3 KR Domain Classification

Sequences were classified based on the subtype motifs of the KR domains. The ones with LDD motifs were labelled as type B KR domains, the ones with a tryptophan eight residues before the catalytic tyrosine were labelled as type A. Further classification was made based on the three residues before the catalytic tyrosine. Since leucine, histidine, and glutamine residues are conserved in B2 type, A2 type and A1-B1 types of KR domains, respectively [39], a further classification was made based on these specifications. KRs that did not bear any of those sequence motifs were labelled as ‘Unclassified’ and the ones that have both patterns of A and B types are labelled as ‘KRbothmotifs’. These groups are not shown in the sequence-position mapping plots. Among all the sequences only 81 of them do not have KR domains. The most abundant type in the alignment is type-B1, with 763 sequences. There are also 6 type-B2, 239 type-A1, 73 type-A2 and 67 type-C KR present. Unfortunately, 996 sequences could not be classified and 20 of them bear motifs of both type A and type B KR domains.

### 4.4 Similarity analysis

The similarity score (ss) was calculated as

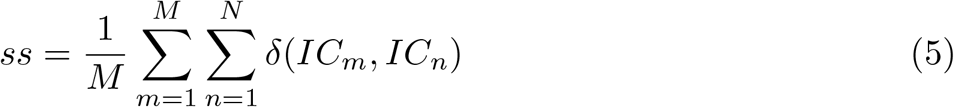

where *δ*(*IC_m_, IC_n_*) is equal to one if the ratio of the number of mutual residues between two ICs to the number of residues in the IC with fewer residues (min(len(*IC_m_*),len(*IC_n_*))) is greater than a threshold (set at 0.7) and either *IC_m_* or *IC_n_* is not matched with another IC; zero otherwise.

### 4.5 Determination of coevolved residue groups

In order to detect the coevolved residue groups, the first step is to calculate the covariation of the amino acids along the MSA:

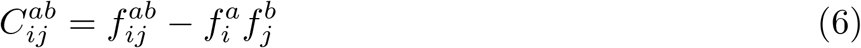

where 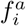 is the frequency of the amino acid *a* at position *i*; 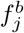 is the frequency of amino acid *b* at position *j*. 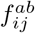 is their joint probability, which measures the probability of *a* and *b* being at positions *i* and *j* simultaneously.

Position-specific conservation is calculated by the Kullback-Leibler relative entropy and the covariance matrix 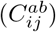 is weighted by the conservation score of each amino acid at each position with respect to the amino acid frequency, which is denoted by 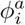 for amino acid *a* at position *i* and 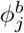 for amino acid *b* at position *j*. This gives the conservation weighted correlation matrix, 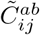, where:

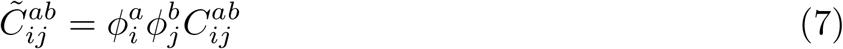

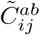, is a four-dimensional matrix (LxLx20×20, where L is the length of the sequence) and it is compressed to a two-dimensional LxL matrix by taking the Frobenius norm. The resulting matrix 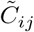 (or statistical coupling analysis (SCA) matrix) is used to determine the coevolved residue groups.

The matrix 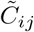 gives the couplings between the positions of the amino acids but doesn’t define networks of co-evolving residues that might have specific subfunction such as catalytic activity, ligand binding, or allosteric signalling. To do that, we need to transform 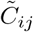 in a way that can separate the residues as different groups and these residue groups should be as independent as possible from each other for the proper detection of a residue group-function relation. For this transformation, firstly, eigenvalue (spectral) decomposition is applied to the 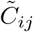. This process decomposes 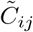 into three matrices:

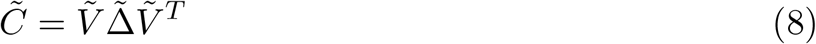

where 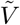 is an eigenvector matrix and 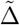 is an eigenvalue matrix, which is a diagonal matrix bearing the weight for each corresponding eigenvector. The matrix 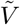 carries the scores that indicate how to combine the residue positions to form the eigenvector. 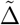 contains the eigenvalues of 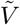 that carries the information about the “importance” of the corresponding eigenvector. For example, a low eigenvalue means an insignificant contribution of the corresponding eigenvector to the variance of the dataset whereas a high eigenvalue means a strong contribution of the corresponding eigenvector to the variance in 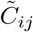. Therefore, the eigenvectors with high eigenvalues carry the information of a “strong” relationship between the residue positions. This means these eigenvectors carry the information of residue positions that have a coevolutionary relationship. On the other hand, the eigenvectors define vectors defining the directions of variance, not the vectors that optimally separate the residues into statistically independent groupings. Indeed, there can be strong couplings between the residues contributing to different eigenvectors. To minimize this interaction between the groups, the significant eigenvectors (the eigenvectors with high weights, 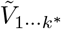) are transformed into new components, which are maximally independent from each other.

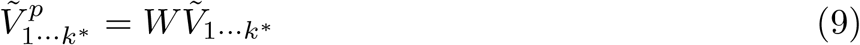

where *W* is a transformation matrix. This mathematical transformation is known as independent component analysis (ICA) [62, 63] and resulting residue groups are called independent components (ICs) [34].

40 independent components are detected; however, only 34 of them have at least two residues in the groups. Therefore, six of them were not taken into account for the further analyses.

When the first eigenvalue is much larger than the remaining significant eigenvalues, it indicates that the first eigenvector is the “coherent” mode and it describes the contribution to all positions to the total correlation [33]. In our system, the first eigenvalue is 822, whereas the second highest is only 143, indicating the dominant first mode. As the first mode has the contributions from all positions, it was removed for the SCA matrix visualization and hierarchical clustering for a clearer detection of the covariation of the residue groups.

### 4.6 Hierarchical clustering of ICs

We performed hierarchical clustering based on average coupling scores of inter-ICs. In the coupling matrix 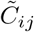, the average coupling score of each square (scores of intra-ICs and inter-ICs) is calculated ending up with 34 x 34 size matrix (for the analysis after removal of some ICs, the size of the average-scored coupling matrix is 22 x 22). Hierarchical clustering was applied on the average-coupling score matrix by scipy.cluster hierarchical clustering package where complete linkage calculations were performed on the average-coupling matrix as distance matrix and the clusters were generated from the calculated linkage matrix.

### 4.7 Sequence-position mapping

Covariation of the MSA matrix columns reveals the coupling between pairs of residue positions and ultimately the detection groups of co-evolving residues (i.e ICs). Similarly, covariation of the MSA matrix rows is expected to give information about groups of related sequences. I.e. the sequences, which have a high correlation between the rows of the MSA matrix, are the sequences with high amino acid sequence similarity.

Since both sequence correlations and position correlations are obtained from rows and columns (respectively) of the same matrix (MSA), they are related to each other by a mathematical approach known as singular value decomposition (SVD).

Basically, SVD is used to decompose any matrix into three matrices:

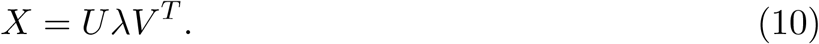

Here, *U* carries the information about sequence similarities (rows-based analysis) and *V* carries information about position similarities (columns-based analysis). *λ* is a diagonal matrix that carries information about which parts of the *U* and *V* matrices provide the “more important” contributions to the *X* matrix.

With a slight modification, the matrix *U*, which carries the sequence similarity pattern, can be obtained from the alignment matrix, *X*, and the position correlation matrix, *V*:

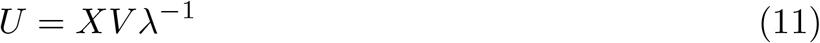

This means that we can obtain sequence similarity patterns specific to coevolved residue groups i.e. independent components. However, since we applied modifications on the positional correlations of the *X* matrix to obtain the coupling matrix 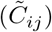, we need to follow the same transformations.

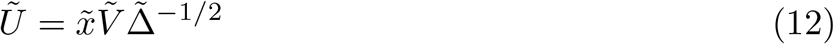

where 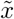 is a compressed alignment matrix (from MxLx20 to MxL), 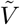 and 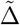 are eigenvectors and eigenvalues of 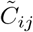, respectively. And as the final step, the ICA transformation is applied:

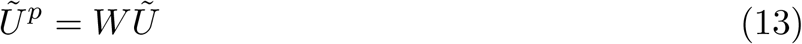

where *W* is the same transformation matrix that was used to transform the top eigenvectors to independent components.

The final 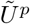 matrix carries the information of sequence divergence patterns of the coevolved residue groups (independent components). Therefore, this analysis allows us to map positional correlations onto sequences and thus this protocol is referred to as sequence-position mapping. For a more detailed explanation of the method please see [34].

### 4.8 Sequence clustering analysis

Thr amino acids contributing to the ICs (ICs 1, 4, 6, 8 and 17) and the sequences of the domains (AT and KR) were clustered using ClustalW [64]. Clustering was performed using the neighbour-joining method (default setting). For visualization of the tree the ETE Toolkit [65] was used.

## 5 Supporting Information

**S1 Fig. The Query Sequence Used for Constructing the MSA: DEBS module 1 plus KS2.**

**S2 Fig. As the number of sequences in the subsampled MSAs increases the similarity score converges to 1.** 100 sequences in the subsampled MSAs are generally insufficient to achieve the same set of ICs as the source MSA. Increasing the number of sequences in the subsampled MSAs eventually gives a saturation point (Ms), the moment at which the similarity score stabilises as 1, which varies among the proteins.

**S3 Fig. Some ICs can be detected successfully even when few sequences are used in the alignment.** A light gray star indicates one occurance in three replicas, a gray star indicates two occurances in three replicas and a black star indicates three occurances in three replicas.

**S4 Fig. The reproduciblity of an IC is independent of the length of the IC.** The number of residues in an IC is plotted against the number of times the IC was detected in three samplings of 100 sequences from the MSA, i.e. when *M′* =100.

**S5 Fig. The similarity of ICs from subsampled MSAs to those from an MSA of 682 sequences of domain composition KS1_AT_KR_ACP_KS2 only.** As the number of sequences in the sub-sampled MSA increases so does the similarity of the ICs to that of the full alignment. The similarity score from subsampled MSAs is rises to a higher value.

**S6 Fig. The similarity of ICs from subsampled MSAs to those from an MSA of 2245 sequences with sequence similarity to the DEBS1 (KS1_AT_KR_ACP_KS2) sequence but varying domain composition.** As the number of sequences in the sub-sampled MSA increases so does the similarity of the ICs to that of the full alignment. Increasing the number of sequences in the MSA to 2245 improves the agreement between the subsampled alignements and the full alignment, as compared to the data from the full alignment with 668 sequences (previous figure).

**S7 Fig. SCA/sectors analysis revealed 34 coevolved residue groups (ICs) spanning one domain or multiple domains and linkers, which can be clustered based on some cross-coupling between the ICs.** The clustering analysis (left side of the figure) was based on the average coupling score between the ICs. The distributions of the residues on each coevolved residue groups (independent component, IC) along the target sequence are shown. KS: Ketosynthase, AT: Acyltransferase, KAL: KS-AT linker, PAL1: post-AT linker 1, PAL2c: post-AT linker 2 conserved region, PAL2nc: post-AT linker 2 non-conserved region, KRs: Ketoreductase structural integrity region, KRc: Ketoreductase catalytic region, ACP: acyl carrier protein, KS2: ketosynthase of module 2.

**S8 Fig. Some ICs can be detected consistently even though the number of sequences in the alignments vary.** ICs, which were detected fewer than or equal to three times in the overall bootstrapping analysis (ICs 9, 19, 20, 23, 24, 28, 29, 31, 32, 33, 34), and the one with two residues (IC_25) were removed from further analysis, resulting in 22 ICs of principle focus. A light gray star indicates one occurance in three replicas, a gray star indicates two occurances in three replicas and a black star indicates three occurances in three replicas.

**S9 Fig. Inter and intra-coupling scores of 34 independent components (ICs) detected via SCA/sector analysis.** The residues on the plot are grouped by IC so the axes cannot be read as amino acid sequential order. Some of the axes labels sit out of line with the other labels due to the very few members in that IC leading to too little space for the label. The colour key represents the scores from the SCA reconstructed without the first eigenvalue and eigenvector.

**S10 Fig. ICs found using an MSA with sequences that must include a KR domain provided domain boundaries consistent with the analysis of the MSA with all sequences similar to the DEBS1 search term.** The hierarchical clustering of the ICs is shown on the left of each panel. The residues in the upper panel are indicated only if they pass the threshold test for inclusion in a given IC, with each IC assigned its own colour. The residues in the lower panel are shaded according to their contribution to the IC, dark indicating a strong contribution, with values shown for all the residues for all ICs (i.e. no statistical test for inclusion has been applied in this panel). KS1:Ketosynthase of module 1, AT: Acyltransferase, KAL: KS-AT linker, PAL1: post-AT linker 1, PAL2c:post-AT linker 2 conserved region, PAL2nc: post-AT linker 2 non-conserved region, KRs: Ketoreductase structural region, KRc: Ketoreductase catalytic region, ACP: Acyl carrier protein, KS2: Ketosynthase of module 2.

**S11 Fig. Logos of the ICs with highly conserved amino acid positions, which cluster together in the hierarchical clustering.** Sequence logos are shown for IC_2, IC_3, IC_5, IC_7, IC_11, IC_13.

**S12 Fig. The average conservation score (Kullback-Leibler relative entropy) of the ICs.** The average conservation score of an IC was determined by taking the average of the Kullback-Leibler relative entropy of all the positions in corresponding IC. The average conservation scores of ICs 2, 3, 5, 7, 11 and 13 are high.

**S13 Fig. Highlighting the variation of amino acid type in the MSA columns associated with IC_2 and IC3.** The sequence positions associated with IC2 or IC_3 were extracted from the MSA and filtered by hhfilter, from hhsuite [66], to have only sequences with ¡80% sequence ID with respect to each other, to highlight the diversity of amino acid type at the different positions. These were displayed as logos plots. (A) logos for IC_2. (B) logos for IC_3.

**S14 Fig. High coupling between the catalytic residues of KS, ACP and KR domains was detected.** The couping matrix that only includes the residues detected in ICs 2, 3, 5, 7, 11 and 13 reveals high coupling between these ICs (A). The catalytic residues of the KS, ACP and KR domains are shown with blue, yellow and green stars in the couping matrix (A) and high coupling can be seen when the catalytic residue coupling matrix is extracted (B). The axes are organised by grouping residues from the same IC, so do not represent the sequential order of the amino acids in the protein.

**S15 Fig. Using only *cis*-AT sequences in the MSA provides similar AT domain boundaries to the whole MSA analysis.** The hierarchical clustering of the ICs is shown on the left of each panel. The residues in the upper panel are indicated only if they pass the threshold test for inclusion in a given IC, with each IC assigned its own colour. The residues in the lower panel are shaded according to their contribution to the IC, dark indicating a strong contribution to the IC, with values shown for all the residues for all ICs (i.e. no statistical test for inclusion has been applied in this panel). KS1:Ketosynthase of module 1, AT: Acyltransferase, KAL: KS-AT linker, PAL1: post-AT linker 1, PAL2c: post-AT linker 2 conserved region, PAL2nc: post-AT linker 2 non-conserved region, KRs: Ketoreductase structural region, KRc: Ketoreductase catalytic region, ACP: Acyl carrier protein, KS2: Ketosynthase of module 2.

**S16 Fig. Sequence-position mapping based on domain compositions of the sequences in the MSA.** In these graphs, if any two class of domain compositions are non-overlapping along an IC, it means the amino acid patterns within the IC are distinct between these compositions. Although for most of the ICs, there is no clear distinction, IC_4, IC_6, IC_8, and IC_17 show separation of some domain compositions.

**S17 Fig. Sequence-position mapping based on extender unit specificity of the AT domains.** Separation of AT domains with different extender unit specificities is seen along ICs 1, 4, 6 and 8.

**S18 Fig. Sequence clustering analysis related to the AT domain.** Sequence clustering analysis of the whole AT domain sequences (A), positions of IC_4 (B) and positions of IC_8 (C) are shown. Different groups of sequences are presented with different colors; blue represents *trans-*AT sequences, red represents malonyl specific AT sequences, purple represents methylmalonyl specific AT sequences, cyan represents ethylmalonyl specific AT sequences, green represents sequences which could not be classified.

**S19 Fig. Residues 760 and 763, contributing strongly to the separation of methylmalonyl and malonyl specific AT domains along IC4, are not highly conserved but have high coupling with residue 753 and are close in the protein structure.** (A) The 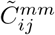 matrix reveals that positions 760 and 763 have strong coupling with position 753. (B) On a 3D model of the structure of the AT from DEBS1 (provided by Louis Woodhouse/Ruairi O’Brien), residues 753, 760 and 763 are close to each other. Highly abundant pairs for positions 760 and 753 in methylmalonyl-specific sequences are Trp-Phe, Leu-Tyr and Arg-Tyr (C); whereas a similar pattern is not observed for malonyl-specific sequences (D).

**S20 Fig. Sequence-position mapping based on subtypes of the KR domain.** Although for most of the ICs, there is no clear distinction in the sequence patterns of the different KR specificities, IC_17 shows some separation of subtypes.

**S21 Fig. SCA analysis of the MSA generated based on the stepwise alignment of sequences with the same domain composition revealed ICs similar to the ICs detected with the original alignment protocol MSA.**

**S1 Table. Comparison of ICs from the full MSA and ICs from the MSA with sequences required to have a KR domain.** Pairs of ICs are shown only if their similarity is higher than 0.7. ICs important for determination of the KR domain boundary are shown in bold.

**S2 Table. Comparison of ICs from the full MSA and ICs from the MSA with only *cis*-AT sequences.** Pairs of ICs are shown only if their similarity is higher than 0.7. ICs important for determination of the AT domain boundary are shown in bold.

**S3 Table. Comparison of the ICs of from the original full MSA and the MSA generated based on the stepwise alignment of sequences with the same domain composition.** Comparison is shown if the IC similarity is higher than 0.6. ICs important for the determination of domain boundaries or with distinct amino acid patterns in different domain sub-types are shown bold.

**S4 Table. Comparison of ICs that determine the domain boundaries and the residues detected as functional in domain subtype specificity between the full MSA and the MSA generated based on domain composition.**

## Acknowledgments

We would like to thank to University of Birmingham for providing the access to BlueBEAR HPC service and covering open access publishing costs.

## References

1. Newman DJ, Cragg GM. Natural Products As Sources of New Drugs over the 30 Years from 1981 to 2010. Journal of Natural Products. 2012;75(3):311–335. doi:10.1021/np200906s.

2. Chan YA, Podevels AM, Kevany BM, Thomas MG. Biosynthesis of polyketide synthase extender units. Nat Prod Rep. 2009;26(1):90–114. doi:10.1039/b801658p.

3. Dutta S, Whicher JR, Hansen DA, Hale WA, Chemler JA, Congdon GR, et al. Structure of a modular polyketide synthase. Nature. 2014;510(7506):512–517. doi:10.1038/nature13423.

4. Robbins T, Liu YC, Cane DE, Khosla C. Structure and mechanism of assembly line polyketide synthases. Current Opinion in Structural Biology. 2016;41:10–18. doi:10.1016/j.sbi.2016.05.009.

5. Keatinge-Clay AT. Polyketide Synthase Modules Redefined. Angewandte Chemie International Edition. 2017;56(17):4658–4660. doi:10.1002/anie.201701281.

6. Vander Wood DA, Keatinge-Clay AT. The modules of trans -acyltransferase assembly lines redefined with a central acyl carrier protein. Proteins: Structure, Function, and Bioinformatics. 2018;86(6):664–675. doi:10.1002/prot.25493.

7. Barajas JF, Blake-Hedges JM, Bailey CB, Curran S, Keasling JD. Engineered polyketides: Synergy between protein and host level engineering. Synthetic and Systems Biotechnology. 2017;2(3):147 – 166. doi:https://doi.org/10.1016/j.synbio.2017.08.005.

8. Cane DE, Liang TC, Taylor PB, Chang C, Yang CC. Macrolide biosynthesis. 3. Stereochemistry of the chain-elongation steps of erythromycin biosynthesis. Journal of the American Chemical Society. 1986;108(16):4957–4964. doi:10.1021/ja00276a042.

9. Keatinge-Clay AT. A Tylosin Ketoreductase Reveals How Chirality Is Determined in Polyketides. Chemistry & Biology. 2007;14(8):898 – 908. doi:https://doi.org/10.1016/j.chembiol.2007.07.009.

10. Weissman KJ. Genetic engineering of modular PKSs: from combinatorial biosynthesis to synthetic biology. Nat Prod Rep. 2016;33:203–230. doi:10.1039/C5NP00109A.

11. Musiol-Kroll EM, Wohlleben W. Acyltransferases as Tools for Polyketide Synthase Engineering. Antibiotics. 2018;7(3). doi:10.3390/antibiotics7030062.

12. Kornfuehrer T, Eustáquio AS. Diversification of polyketide structures via synthase engineering. Med Chem Commun. 2019; p. –. doi:10.1039/C9MD00141G.

13. Beck C, Garzón JFG, Weber T. Recent Advances in Re-engineering Modular PKS and NRPS Assembly Lines. Biotechnology and Bioprocess Engineering. 2020;25(6):886–894. doi:10.1007/s12257-020-0265-5.

14. Oliynyk M, Brown MJB, Cortés J, Staunton J, Leadlay PF. A hybrid modular polyketide synthase obtained by domain swapping. Chemistry & Biology. 1996;3(10):833 – 839. doi:https://doi.org/10.1016/S1074-5521(96)90069-1.

15. Ruan X, Pereda A, Stassi DL, Zeidner D, Summers RG, Jackson M, et al. Acyltransferase domain substitutions in erythromycin polyketide synthase yield novel erythromycin derivatives. Journal of Bacteriology. 1997;179(20):6416–6425. doi:10.1128/jb.179.20.6416-6425.1997.

16. Liu L, Thamchaipenet A, Fu H, Betlach M, Ashley G. Biosynthesis of 2-Nor-6-deoxyerythronolide B by Rationally Designed Domain Substitution. Journal of the American Chemical Society. 1997;119(43):10553–10554. doi:10.1021/ja972451y.

17. Lau J, Fu H, Cane DE, Khosla C. Dissecting the Role of Acyltransferase Domains of Modular Polyketide Synthases in the Choice and Stereochemical Fate of Extender Units. Biochemistry. 1999;38(5):1643–1651. doi:10.1021/bi9820311.

18. Petkovic H, Lill RE, Sheridan RM, Wilkinson B, McCormick EL, McArthur HAI, et al. A Novel Erythromycin, 6-Desmethyl Erythromycin D, Made by Substituting an Acyltransferase Domain of the Erythromycin Polyketide Synthase. The Journal of Antibiotics. 2003;56(6):543–551. doi:10.7164/antibiotics.56.543.

19. Yuzawa S, Deng K, Wang G, Baidoo EEK, Northen TR, Adams PD, et al. Comprehensive in Vitro Analysis of Acyltransferase Domain Exchanges in Modular Polyketide Synthases and Its Application for Short-Chain Ketone Production. ACS Synthetic Biology. 2017;6(1):139–147. doi:10.1021/acssynbio.6b00176.

20. Kellenberger L, Galloway IS, Sauter G, Böhm G, Hanefeld U, Cortés J, et al. A Polylinker Approach to Reductive Loop Swaps in Modular Polyketide Synthases. ChemBioChem;9(16):2740–2749. doi:10.1002/cbic.200800332.

21. Valenzano CR, Lawson RJ, Chen AY, Khosla C, Cane DE. The Biochemical Basis for Stereochemical Control in Polyketide Biosynthesis. Journal of the American Chemical Society. 2009;131(51):18501–18511. doi:10.1021/ja908296m.

22. Annaval T, Paris C, Leadlay PF, Jacob C, Weissman KJ. Evaluating Ketoreductase Exchanges as a Means of Rationally Altering Polyketide Stereochemistry. ChemBioChem;16(9):1357–1364. doi:10.1002/cbic.201500113.

23. Eng CH, Yuzawa S, Wang G, Baidoo EEK, Katz L, Keasling JD. Alteration of Polyketide Stereochemistry from anti to syn by a Ketoreductase Domain Exchange in a Type I Modular Polyketide Synthase Subunit. Biochemistry. 2016;55(12):1677–1680. doi:10.1021/acs.biochem.6b00129.

24. Reeves CD, Murli S, Ashley GW, Piagentini M, Hutchinson CR, McDaniel R. Alteration of the Substrate Specificity of a Modular Polyketide Synthase Acyltransferase Domain through Site-Specific Mutations. Biochemistry. 2001;40(51):15464–15470. doi:10.1021/bi015864r.

25. Del Vecchio F, Petkovic H, Kendrew SG, Low L, Wilkinson B, Lill R, et al. Active-site residue, domain and module swaps in modular polyketide synthases. Journal of Industrial Microbiology and Biotechnology. 2003;30(8):489–494. doi:10.1007/s10295-003-0062-0.

26. Sundermann U, Bravo-Rodriguez K, Klopries S, Kushnir S, Gomez H, Sanchez-Garcia E, et al. Enzyme-Directed Mutasynthesis: A Combined Experimental and Theoretical Approach to Substrate Recognition of a Polyketide Synthase. ACS Chemical Biology. 2013;8(2):443–450. doi:10.1021/cb300505w.

27. Zhang F, Shi T, Ji H, Ali I, Huang S, Deng Z, et al. Structural Insights into the Substrate Specificity of Acyltransferases from Salinomycin Polyketide Synthase. Biochemistry. 2019;58(27):2978–2986. doi:10.1021/acs.biochem.9b00305.

28. Zheng J, Piasecki SK, Keatinge-Clay AT. Structural Studies of an A2-Type Modular Polyketide Synthase Ketoreductase Reveal Features Controlling *α*-Substituent Stereochemistry. ACS Chemical Biology. 2013;8(9):1964–1971. doi:10.1021/cb400161g.

29. Bailey CB, Pasman ME, Keatinge-Clay AT. Substrate structure–activity relationships guide rational engineering of modular polyketide synthase ketoreductases. Chem Commun. 2016;52:792–795. doi:10.1039/C5CC07315D.

30. Korber BT, Farber RM, Wolpert DH, Lapedes AS. Covariation of mutations in the V3 loop of human immunodeficiency virus type 1 envelope protein: an information theoretic analysis. Proceedings of the National Academy of Sciences. 1993;90(15):7176–7180. doi:10.1073/pnas.90.15.7176.

31. Morcos F, Pagnani A, Lunt B, Bertolino A, Marks DS, Sander C, et al. Direct-coupling analysis of residue coevolution captures native contacts across many protein families. Proceedings of the National Academy of Sciences. 2011;108(49):E1293–E1301. doi:10.1073/pnas.1111471108.

32. Ekeberg M, Lövkvist C, Lan Y, Weigt M, Aurell E. Improved contact prediction in proteins: Using pseudolikelihoods to infer Potts models. Phys Rev E. 2013;87:012707. doi:10.1103/PhysRevE.87.012707.

33. Halabi N, Rivoire O, Leibler S, Ranganathan R. Protein Sectors: Evolutionary Units of Three-Dimensional Structure. Cell. 2009;138(4):774–786. doi:10.1016/j.cell.2009.07.038.

34. Rivoire O, Reynolds KA, Ranganathan R. Evolution-Based Functional Decomposition of Proteins. PLOS Computational Biology. 2016;12(6):1–26. doi:10.1371/journal.pcbi.1004817.

35. Narayanan C, Gagné D, Reynolds KA, Doucet N. Conserved amino acid networks modulate discrete functional properties in an enzyme superfamily. Scientific Reports. 2017;7(1):3207. doi:10.1038/s41598-017-03298-4.

36. Scheibenreif L, Littmann M, Orengo C, Rost B. FunFam protein families improve residue level molecular function prediction. BMC Bioinformatics. 2019;20(1). doi:10.1186/s12859-019-2988-x.

37. Das S, Scholes HM, Sen N, Orengo C. CATH functional families predict functional sites in proteins. Bioinformatics. 2020;37(8):1099–1106. doi:10.1093/bioinformatics/btaa937.

38. Keatinge-Clay AT, Stroud RM. The Structure of a Ketoreductase Determines the Organization of the *β*-Carbon Processing Enzymes of Modular Polyketide Synthases. Structure. 2006;14(4):737 – 748. doi:https://doi.org/10.1016/j.str.2006.01.009.

39. Zheng J, Keatinge-Clay AT. The status of type I polyketide synthase ketoreductases. Med Chem Commun. 2013;4:34–40. doi:10.1039/C2MD20191G.

40. Kautsar SA, Blin K, Shaw S, Navarro-Muñoz JC, Terlouw BR, van der Hooft JJJ, et al. MIBiG 2.0: a repository for biosynthetic gene clusters of known function. Nucleic Acids Research. 2019;doi:10.1093/nar/gkz882.

41. Oefner C, Schulz H, D Arcy A, Dale GE. Mapping the active site ofEscherichia colimalonyl-CoA–acyl carrier protein transacylase (FabD) by protein crystallography. Acta Crystallographica Section D Biological Crystallography. 2006;62(6):613–618. doi:10.1107/s0907444906009474.

42. Smith S, Tsai SC. The type I fatty acid and polyketide synthases: a tale of two megasynthases. Natural Product Reports. 2007;24(5):1041. doi:10.1039/b603600g.

43. Haydock SF, Aparicio JF, Molnár I, Schwecke T, Khaw LE, König A, et al. Divergent sequence motifs correlated with the substrate specificity of (methyl)malonyl-CoA:acyl carrier protein transacylase domains in modular polyketide synthases. FEBS Letters. 1995;374(2):246–248. doi:10.1016/0014-5793(95)01119-y.

44. Russ WP, Lowery DM, Mishra P, Yaffe MB, Ranganathan R. Natural-like function in artificial WW domains. Nature. 2005;437(7058):579–583. doi:10.1038/nature03990.

45. Reynolds KA, McLaughlin RN, Ranganathan R. Hot Spots for Allosteric Regulation on Protein Surfaces. Cell. 2011;147(7):1564–1575. doi:10.1016/j.cell.2011.10.049.

46. Weigt M, White RA, Szurmant H, Hoch JA, Hwa T. Identification of direct residue contacts in protein-protein interaction by message passing. Proceedings of the National Academy of Sciences. 2008;106(1):67–72. doi:10.1073/pnas.0805923106.

47. Morcos F, Pagnani A, Lunt B, Bertolino A, Marks DS, Sander C, et al. Direct-coupling analysis of residue coevolution captures native contacts across many protein families. Proceedings of the National Academy of Sciences. 2011;108(49):E1293–E1301. doi:10.1073/pnas.1111471108.

48. Devlin J, Chang MW, Lee K, Toutanova K. BERT: Pre-training of Deep Bidirectional Transformers for Language Understanding; 2019.

49. Michel M, Hurtado DM, Elofsson A. PconsC4: fast, accurate and hassle-free contact predictions. Bioinformatics. 2018;35(15):2677–2679. doi:10.1093/bioinformatics/bty1036.

50. Ji S, Oruç T, Mead L, Rehman MF, Thomas CM, Butterworth S, et al. DeepCDpred: Inter-residue distance and contact prediction for improved prediction of protein structure. PLOS ONE. 2019;14(1):e0205214. doi:10.1371/journal.pone.0205214.

51. Xu J. Distance-based protein folding powered by deep learning. Proceedings of the National Academy of Sciences. 2019;116(34):16856–16865. doi:10.1073/pnas.1821309116.

52. Senior AW, Evans R, Jumper J, Kirkpatrick J, Sifre L, Green T, et al. Improved protein structure prediction using potentials from deep learning. Nature. 2020;577(7792):706–710. doi:10.1038/s41586-019-1923-7.

53. Mirabello C, Wallner B. rawMSA: End-to-end Deep Learning using raw Multiple Sequence Alignments. PLOS ONE. 2019;14(8):e0220182. doi:10.1371/journal.pone.0220182.

54. Sercu T, Verkuil R, Meier J, Amos B, Lin Z, Chen C, et al. Neural Potts Model. bioRxiv. 2021;doi:10.1101/2021.04.08.439084.

55. Tondnevis F, Dudenhausen EE, Miller AM, McKenna R, Altschul SF, Bloom LB, et al. Deep Analysis of Residue Constraints (DARC): identifying determinants of protein functional specificity. Scientific Reports. 2020;10(1). doi:10.1038/s41598-019-55118-6.

56. Wang SW, Bitbol AF, Wingreen NS. Revealing evolutionary constraints on proteins through sequence analysis. PLOS Computational Biology. 2019;15(4):e1007010. doi:10.1371/journal.pcbi.1007010.

57. Rives A, Meier J, Sercu T, Goyal S, Lin Z, Liu J, et al. Biological structure and function emerge from scaling unsupervised learning to 250 million protein sequences. Proceedings of the National Academy of Sciences. 2021;118(15):e2016239118. doi:10.1073/pnas.2016239118.

58. Gouldson PR, Winn PJ, Reynolds CA. A Molecular Dynamics Approach to Receptor Mapping: Application to the 5HT3 and .beta.2-Adrenergic Receptors. Journal of Medicinal Chemistry. 1995;38(20):4080–4086. doi:10.1021/jm00020a023.

59. Remmert M, Biegert A, Hauser A, Söding J. HHblits: lightning-fast iterative protein sequence searching by HMM-HMM alignment. Nature Methods. 2011;9:173 EP –.

60. El-Gebali S, Mistry J, Bateman A, Eddy SR, Luciani A, Potter SC, et al. The Pfam protein families database in 2019. Nucleic Acids Research. 2018;47(D1):D427–D432. doi:10.1093/nar/gky995.

61. HMMER: biosequence analysis using profile hidden Markov models;. http://hmmer.org/.

62. Bell AJ, Sejnowski TJ. An Information-Maximization Approach to Blind Separation and Blind Deconvolution. Neural Computation. 1995;7(6):1129–1159. doi:10.1162/neco.1995.7.6.1129.

63. Hyvärinen A, Karhunen J, Oja E. Independent component analysis. J. Wiley; 2001.

64. Larkin MA, Blackshields G, Brown NP, Chenna R, McGettigan PA, McWilliam H, et al. Clustal W and Clustal X version 2.0. Bioinformatics. 2007;23(21):2947–2948. doi:10.1093/bioinformatics/btm404.

65. Huerta-Cepas J, Serra F, Bork P. ETE 3: Reconstruction, Analysis, and Visualization of Phylogenomic Data. Molecular Biology and Evolution. 2016;33(6):1635–1638. doi:10.1093/molbev/msw046.

66. Söding J. Protein homology detection by HMM–HMM comparison. Bioinformatics. 2004;21(7):951–960. doi:10.1093/bioinformatics/bti125.

